# Single cell transcriptomics reveals chondrocyte differentiation dynamics *in vivo* and *in vitro*

**DOI:** 10.1101/2023.12.20.572425

**Authors:** John E G Lawrence, Steven Woods, Kenny Roberts, Dinithi Sumanaweera, Petra Balogh, Alexander V Predeus, Peng He, Tong Li, Krzysztof Polanski, Elena Prigmore, Elizabeth Tuck, Lira Mamanova, Di Zhou, Simone Webb, Laura Jardine, Xiaoling He, Roger A Barker, Muzlifah Haniffa, Adrienne M Flanagan, Matthew D Young, Sam Behjati, Omer Bayraktar, Susan J Kimber, Sarah A Teichmann

**Affiliations:** Wellcome Sanger Institute, Wellcome Genome Campus, Hinxton, Cambridge CB10 1SA, UK; Department of Trauma and Orthopaedic Surgery, Addenbrookes Hospital, Cambridge CB2 0QQ, UK; Division of Cell Matrix Biology and Regenerative Medicine, School of Biological Sciences, Faculty of Biology, Medicine and Health, University of Manchester, Manchester M13 9PT, UK; Department of Cellular and Molecular Pathology, Royal National Orthopaedic Hospital, Brockley Hill, Stanmore HA7 4LP, UK; European Molecular Biology Laboratory, European Bioinformatics Institute (EMBL-EBI), Wellcome Genome Campus, Cambridge, UK; Biosciences Institute, Newcastle University, Newcastle upon Tyne, NE2 4HH, UK; Wellcome-MRC Stem Cell Institute and Department of Clinical Neurosciences, University of Cambridge, Cambridge, CB2 0QQ, UK; Research Department of Pathology, University College London (UCL) Cancer Institute, London WC1E 6DD, UK; Department of Paediatrics, University of Cambridge, Cambridge CB2 0QQ, UK

**Keywords:** Cartilage, tissue engineering, single-cell RNA sequencing, skeletal development, in vitro chondrogenesis

## Abstract

The consistent production of *in vitro* chondrocytes that faithfully recapitulate *in vivo* development would be of great benefit for musculoskeletal disease modelling and regenerative medicine. Current efforts are often limited by off-target differentiation, resulting in a heterogeneous product. Furthermore, the lack of comparison to human embryonic tissue, precludes detailed evaluation of *in vitro* cells. Here, we perform single-cell RNA sequencing of embryonic long bones dissected from first trimester hind limbs from a range of gestational ages. We combine this with publicly available data to form a detailed atlas of endochondral ossification, which we then use to evaluate a series of published *in vitro* chondrogenesis protocols, finding substantial variability in cell states produced by each. We apply single-nuclear RNA sequencing to one protocol to enable direct comparison between *in vitro* and *in vivo,* and perform trajectory alignment between the two to reveal differentiation dynamics at the single-cell level, shedding new light on off-target differentiation *in vitro*. Using this information, we inhibit the activity of FOXO1, a transcription factor predicted to be active in embryonic bone development and in chondrogenic cells *in vitro*, and increase chondrocyte transcripts *in vitro.* This study therefore presents a new framework for evaluating tissue engineering protocols, using single-cell data from human development to drive improvement and bring the prospect of true engineered cartilage closer to reality.

## Introduction

In recent years, numerous large-scale single-cell profiling studies have rapidly accelerated our understanding of the cellular composition of multiple organ systems^1^. These efforts extend to human development, with profiles of different systems across developmental time providing insight into the origins of a range of diseases^2,3^. This improved understanding of *in vivo* development promises to improve the fidelity of *in vitro* organoid systems, providing both a benchmark for their evaluation and highlighting points of divergence from human biology to feedback into differentiation protocol development^4,5^. This approach has the broad potential to advance disease modelling, drug development, precision therapy and regenerative medicine.

One field where such advances are urgently required is musculoskeletal biology, as the global burden of disease attributable to this organ system inexorably rises, making it the leading cause of disability worldwide ^6,7^. In particular, the ability to accurately and reliably engineer cartilage *in vitro* would be of great benefit from both a disease modelling and regenerative medicine standpoint ^8–10^. At present, the considerable variability in culture systems and their cell sources together with the propensity of chondrocytes to de-differentiate in culture has made cartilage engineering a challenge^11–14^.

Broadly speaking, present *in vitro* chondrogenesis systems are focused on recapitulating one of two developmental processes; endochondral ossification and articular cartilage formation. In the former, mesenchymal progenitors organise into a condensate before differentiating into immature resting chondrocytes. These subsequently undergo proliferation before entering a transitional phase, known as prehypertrophy, before entering a phase of rapid cellular growth known as hypertrophy. This process of proliferation and hypertrophy, together with the secretion of extracellular matrix, drives bone growth^15^. Surrounding this cartilaginous anlage is a shell of perichondrium, which receives signals from hypertrophic cartilage to differentiate first into periosteum and subsequently into periosteal-derived osteoblasts, which form the “bone collar” around the forming bone, which eventually gives rise to compact, cortical bone^16^. Hypertrophic chondrocytes eventually apoptose, permitting the invasion of blood vessels into the cartilage anlage, which deliver osteoclasts (multinucleate myeloid cells that break down the matrix) and osteoblast precursors, which mineralise the remaining matrix as they mature, giving rise to the inner, trabecular bone^16^. Articular chondrocytes are derived from the interzone, a collection of the aforementioned condensed mesenchymal cells that gather at the sight of the future joint, lose expression of chondrocyte genes such as *COL2A1* and *SOX9* and instead express *GDF5*^17^. These cells act as a common progenitor for all the components of the synovial joints ^18^.

Here, we set out to profile human skeletal development by performing single-cell RNA sequencing on individual long bones dissected from first trimester human hind limbs. We combine this data with that of previous studies to produce a detailed atlas of endochondral ossification^19–21^. We then use this atlas as a reference for evaluating a series of *in vitro* chondrogenesis protocols, finding substantial variability in the cell states produced by the different protocols, all of which exhibited some off-target differentiation. We validate our findings for a protocol predicted to achieve chondrocyte hypertrophy using single-nuclear RNA sequencing and RNA *in situ* hybridisation, applying Genes2Genes alignment to give new insights into chondrocyte differentiation dynamics and quantifying off-target transcription *in vitro*. Finally, we utilise a well characterised inhibitor to block the activity of FOXO1, a transcription factor thought to play a role in bone development, and predicted to be active in fetal osteoblasts and *in vitro* cells in our data ^22^. The FOXO1 inhibitor increased the expression of transcripts characteristic of hypertrophic chondrocytes *in vitro*. Our data can be freely accessed at https://chondrocytes.cellgeni.sanger.ac.uk/

## Results

### Endochondral Ossification at Single Cell Resolution

We performed single-cell RNA sequencing (scRNAseq) on 18 long bone samples dissected from three fetuses aged 7-9 post-conception weeks (PCW) (Fig. 1A). From these, 72,944 cells passed quality control criteria. We then integrated these with data from two previous studies of embryonic limbs (see methods) to produce a developmental atlas of 249,151 cells spanning 5-17 PCW (Fig. 2A & B) Fig. S1A for quality control)^19,20^. Following pre-processing, quality-control and clustering, analysis of classical marker gene expression revealed a diverse set of cell types. These included haematopoetic cells, from both lymphoid (n=13,259) and myeloid compartments (n=35,772, including 261 RANK^+^/CD43^+^ osteoclasts), glia (n=2,681), muscle (n=17,494), dermal (n=2,040), endothelium (n=4,008), tendon (n=6,470), adipose (n=394), osteochondral (n=78,347), fibrous (n=25,153) and mesenchymal progenitors (n=63,533) (Fig. 1A; Fig. 2A & B; supplementary table 1 for marker genes). Within the mesenchymal progenitors category, TBX5^+^ lateral plate mesoderm (LPM) cells were captured, together with two populations of mesenchymal condensate; the first *SOX9*^low^ (Mesenchymal Condensate 1) and the second *SOX9*^high^ (Mesenchymal Condensate 2; Fig. 2A & B). Other progenitors captured included *ISL1*^+^ mesenchyme, two populations of *MEIS*-expressing proximal mesenchyme, and three populations expressing *HOXD13*, denoting cells in the autopod (Distal mesenchyme 1-3; Fig. 2A & B).

**Figure1.**
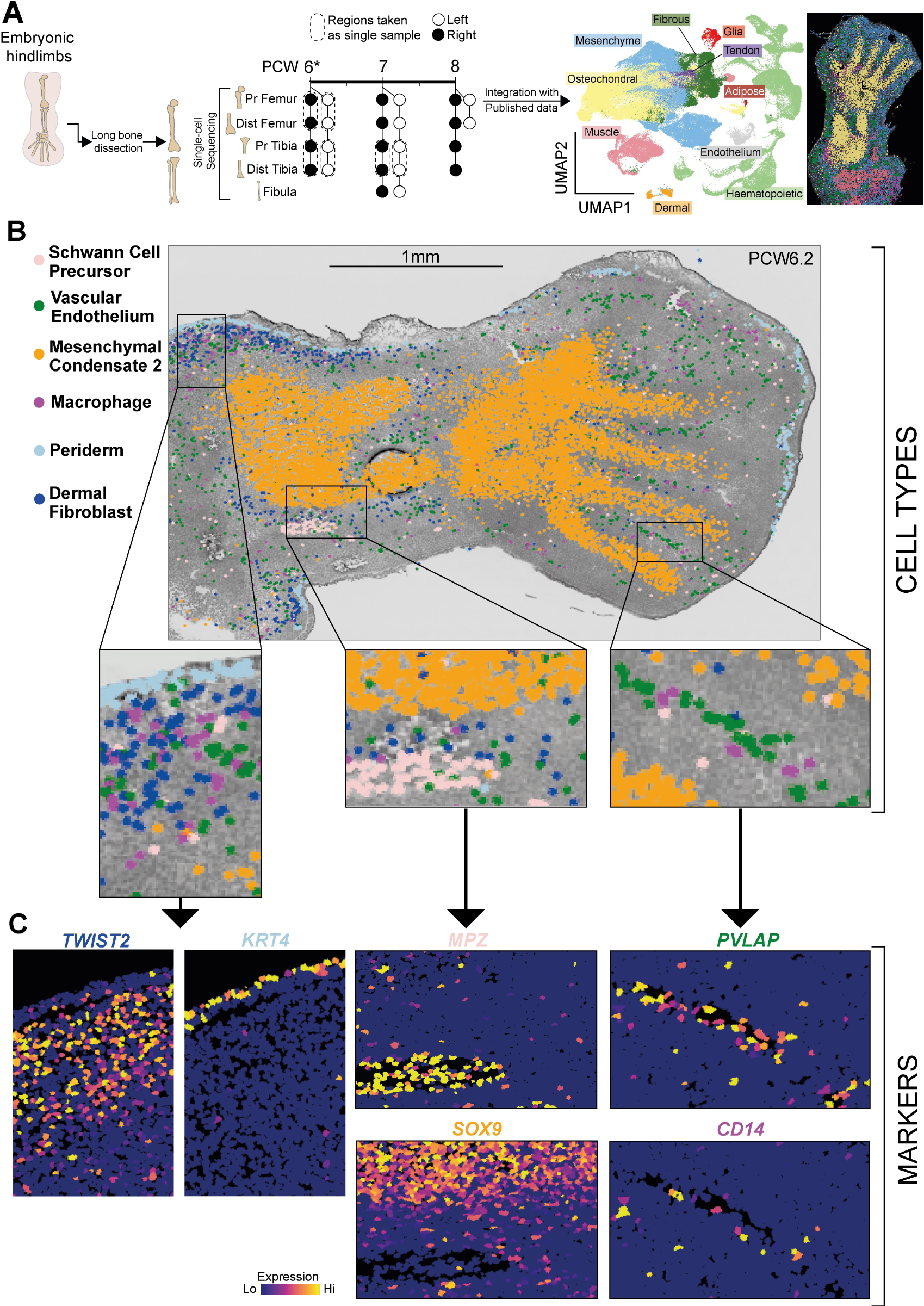
Single-cell level characterisation of the developing appendicular skeleton. **(A)** Left panel-Experimental Overview. PCW denotes post-conception weeks. Pr: proximal, Dist; distal. Right panel-Uniform Manifold Approximation & Projection (UMAP) plot of broad cell categories within the embryonic limb, and broad cell categories mapped onto in-situ sequencing data of the PCW6 hindlimb. **(B)** inferred location of single cells in the PCW6.2 hind limb using k-nearest neighbour prediction on in-situ sequencing data. Coloured points represent predicted cell type location as shown in the legend. Boxed regions represent areas shown at greater magnification in the lower panel. **(C)** Marker gene expression from in-situ sequencing for cell types depicted in (B). Font colour corresponds to cell type marked by that gene. Cells are colour-coded according to gene expression level, ranging from not detected (blue) to the highest detected levels (yellow), according to the adjacent colour key.

**Figure2.**
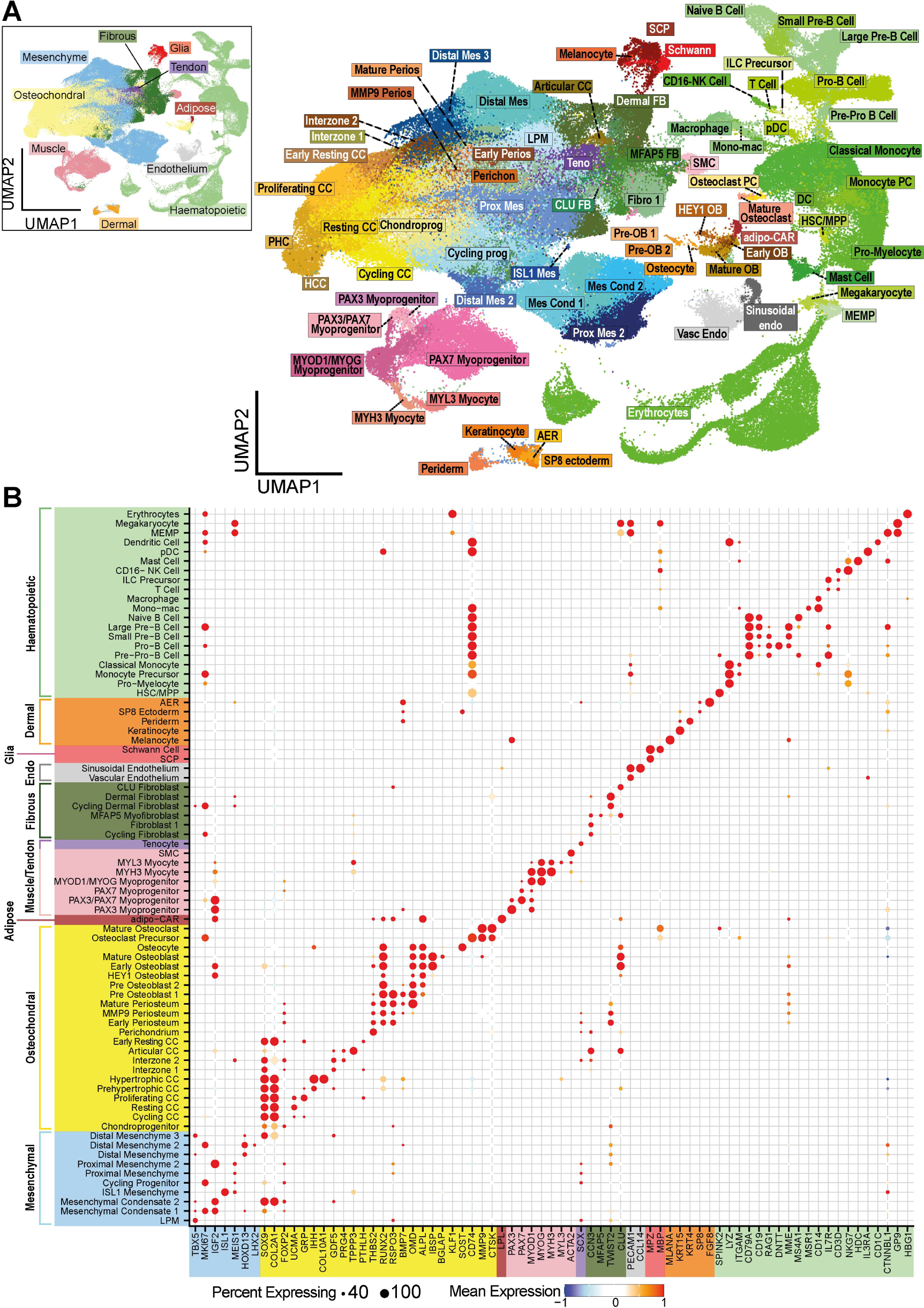
Cell types in the developing human limb. **(A)** Inset panel-Uniform Manifold Approximation & Projection (UMAP) plot of broad cell categories within the embryonic limb. Main / right panel - Uniform Manifold Approximation & Projection (UMAP) plot of finely annotated cell types captured from the embryonic limb. **(B)** Dotplot visualisation showing scaled marker gene expression for cells depicted in (A). Dot size corresponds to the fraction of cells with non-zero expression. Dots are coloured according to gene expression level, according to the adjacent colour key. CC; Chondrocyte, LPM; lateral plate mesoderm, SMC; smooth muscle cell, SCP; schwann cell precursor, AER; apical ectodermal ridge,HSC/MPP; haematopoietic stem cell/ multipotent progenitor, pDC; plasmacytoid dendritic cell, ILC; innate lymphoid cell, MEMP; Megakaryocyte–erythroid progenitor cell.

To place cells in their anatomical context at high resolution, we applied a nearest neighbour technique to combine this integrated single cell atlas with our 90-gene in-situ sequencing (ISS) dataset for the embryonic hindlimb^21^ (see methods). This analysis revealed clear spatial segregation between mesenchymal-derived cell types, such as muscle, cartilage and periosteum, as well as co-location of vascular endothelial cells and macrophages. (Fig. 1B; Fig. S2A). Cells of the periderm and SP8^+^ ectodermal cells were superficial to dermal fibroblasts, and Schwann cell precursors matched to nerve tracts (Fig. 1B; Fig S2A). These predicted locations were confirmed through canonical marker gene expression (Fig. 1C; Fig. S2B).

Focusing on osteochondral tissue, we sub-clustered approximately 55,000 cells in the chondrocyte lineage in order to obtain a transcriptomic readout of the progression of cells through endochondral ossification. This began with chondroprogenitors expressing *PRRX1*, *TWIST1* and *FOXP1*/*2* (Fig. 3A & B)^23–26^. Six clusters of *SOX9*^+^*COL2A1^+^* chondrocytes were also identified. This included a cluster (Early Resting Chondrocytes) expressing *PTHLH*, characteristic of resting chondrocytes, that harboured weak expression of the chondrocyte markers *SOX9*, *COL2A1* and *MATN4*^27^. A second cluster (Resting Chondrocytes) more strongly expressed these three genes, as well as other cartilage extracellular matrix genes including *UCMA*, which is expressed in the distal and peripheral resting zone^28^. Other chondrocytes expressed classical marker genes, including *GRP* (Proliferating), *IHH* (Prehypertrophic) and *COL10A1* (Hypertrophic; Fig. 3A & B)^28–31^. We also identified two clusters of interzone cells; *GDF5*^+^*PRG4*^−^ (Interzone 1) and *GDF5*^+^*PRG4*^+^ (Interzone 2) as well as a cluster of *PRG4*^+^ articular chondrocytes which also expressed *AQP1*, *TPPP3,* and were *COL2A1* negative (Fig. 3A & B)^17,19,32,33^.

**Figure3.**
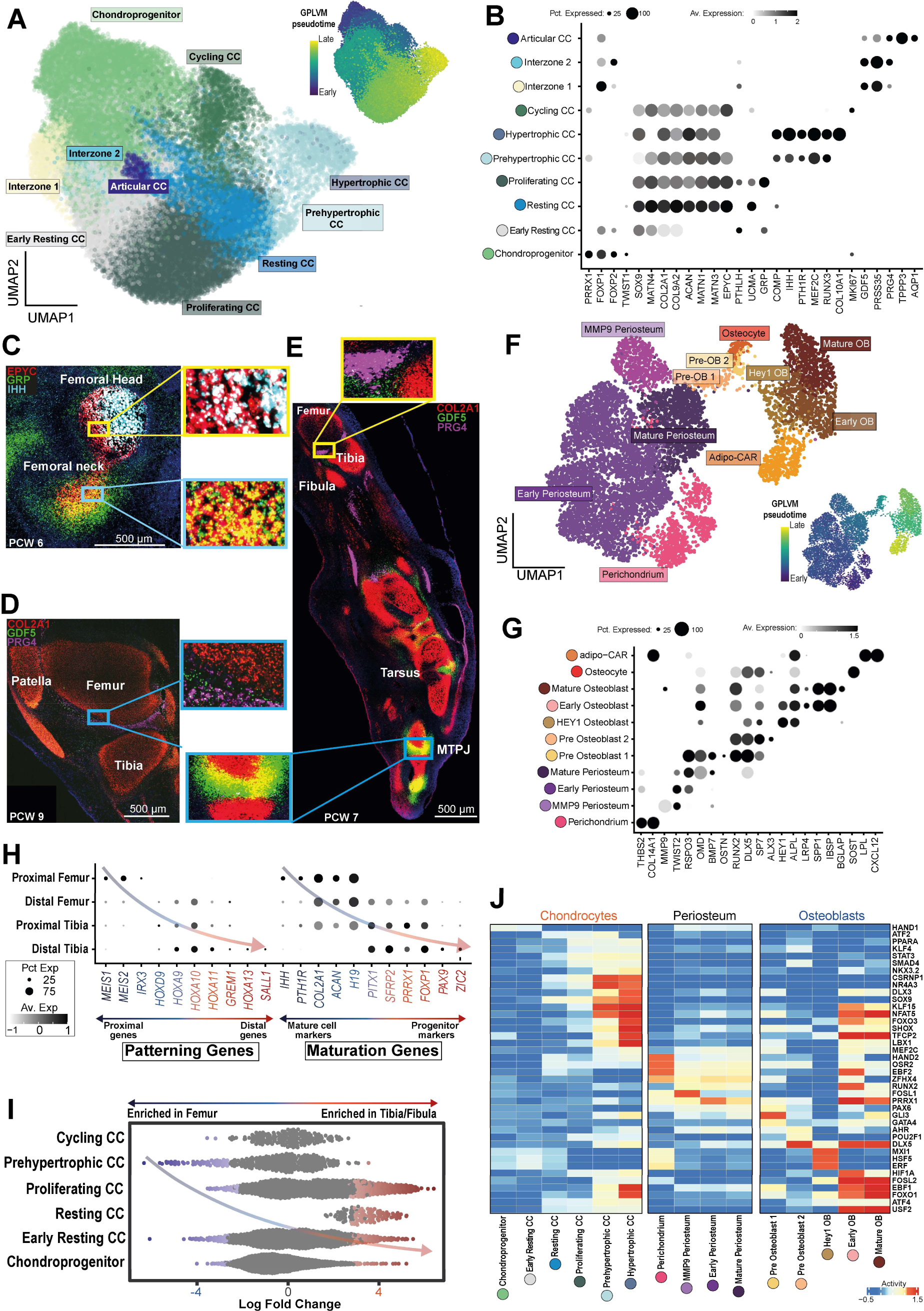
Endochondral ossification in the human embryonic hindlimb. **(A)** Uniform Manifold Approximation & Projection (UMAP) plot showing chondrocyte populations within the developing limb. Inset-GPLVM pseudotime calculation for the same projection, colour-coded according to pseudotime value, ranging from low (blue) to the high(yellow), according to the adjacent colour key **(B)** Dotplot visualisation showing scaled marker gene expression for cells depicted in (A). Dot size corresponds to the fraction of cells with non-zero expression. Dots are coloured according to gene expression level, ranging from not detected (white) to the highest detected levels (black), according to the adjacent colour key. **(C)** RNA-In Situ Hybridisation of the PCW6 femoral head and neck in axial section for *EPYC, GRP* and *IHH*. Inset high magnification boxes represent the regions in the corresponding coloured boxes on the lower magnification image. **(D)** RNA-In Situ Hybridisation of the PCW9 knee joint in sagittal section for *COL2A1*, *GDF5* and *PRG4*. The inset high magnification box represents the region in the corresponding coloured box on the lower magnification image. **(E)** RNA-In Situ Hybridisation of the PCW7 knee joint, zeugopod and autopod in sagittal section for *COL2A1*, *GDF5* and *PRG4*. Inset high magnification boxes represent the regions in the corresponding coloured boxes on the lower magnification image. **(F)** Uniform Manifold Approximation & Projection (UMAP) plot showing periosteal & osteoblast populations within the embryonic limb. Inset-GPLVM pseudotime calculation for the same projection, colour-coded according to pseudotime valuel, ranging from low (blue) to the high(yellow), according to the adjacent colour key. OB; osteoblast. **(G)** Dotplot visualisation showing scaled marker gene expression for cells depicted in (F). Dot size corresponds to the fraction of cells with non-zero expression. Dots are coloured according to gene expression level, ranging from not detected (white) to the highest detected levels (black), according to the adjacent colour key. **(H)** Dotplot visualisation showing scaled gene expression for fetal chondrocytes by sample location. Dot size corresponds to the fraction of cells with non-zero expression. Dots are coloured according to gene expression level, ranging from not detected (white) to the highest detected levels (black), according to the adjacent colour key. **(I)** Beeswarm plot visualising differential abundance of cell types in the femora versus the tibiae and fibulae of the hindlimb. CC; Chondrocyte. **(J)** Heatmap visualising predicted TF activities for cell types depicted in (A) and (F). Colour corresponds to activity level, ranging from low (blue) to high (red), according to the adjacent colour key. CC; chondrocyte, OB; osteoblast

We examined the expression patterns of several of these genes at single cell resolution by applying RNA *in situ* hybridization to hindlimbs aged 6-9 PCW (Fig. 3C-E). This allowed us to visualise the continuum of development along the femoral neck and head at 6 PCW (Fig. 3C). At the knee joint at 9 PCW, *GDF5*^+^ and *PRG4*^+^ cells were visualised in keeping with the two interzone clusters and articular chondrocytes in the single cell data (Fig. 3D). There was a clear transition point between these cells and *COL2A1*^+^ cells of the distal femur and proximal tibia, again reflecting the single cell data (Fig. 3D). By contrast, a sagittal section of the hindlimb at 7 PCW showed *GDF5* upregulation in the digital region outside of the forming synovial joint with numerous *GDF5*^+^/*COL2A1*^+^ cells, in keeping with its role in regulating digit growth (Fig. 3E)^34^.

Our experiments also captured cells from the perichondrium, periosteum and nascent bone collar (Fig. 3F & G). The perichondrium cluster expressed perichondrial-specific genes *THBS2* and *COL14A1*, whilst periosteum clusters expressed the *RUNX2* suppressor *TWIST2* and the osteogenic regulator *RSPO3* (Fig. 3G) ^35–37^. The Mature Periosteum cluster also expressed the pro-osteogenic factors osteomodulin/*OMD* and *BMP7* and had weaker *TWIST2* expression (Fig. 3G)^38,39^. A third periosteal population (MMP9 periosteum) expressed matrix metalloproteinase-9 (*MMP9*; Fig. 3G). This molecule increases the bioavailability of hypertrophic chondrocyte-derived VEGF and stimulates these cells to release MMP13 in order to solubilise cartilage and enable osteoclast and endothelial cell invasion into the developing diaphyseal bone^40^. A further cluster (Pre Osteoblast 1) was similar to periosteal cells, but expressed the pro-osteogenic transcription factors *RUNX2, DLX5* and *SP7/*osterix(Fig.3G). In addition to these genes, the Pre Osteoblast 2 cluster expressed the pro-osteogenic factor *ALX3,* and lacked the periosteal marker gene expression, suggesting further differentiation away from their parent cell type (Fig.3G)^41,42^. These clusters lacked the markers of osteogenic maturation expressed in the three remaining osteoblast clusters, the cells of which originated mostly in the 2nd trimester (Fig. S3A & D), and likely represent periosteal-derived osteoblasts. The final three osteoblast clusters included one expressing the canonical *WNT* target *HEY1* (HEY1 osteoblast), a negative regulator of mineralisation. This cluster expressed *ALPL* but no other markers of maturation (Fig. 3G)^43^. Early osteoblasts expressing *ALPL, SPP1/*osteopontin and *IBSP* were also captured, together with mature cells expressing *BGLAP*/osteocalcin^44^. *SOST/*sclerostin-expressing osteocytes were also captured(Fig. 3 F & G)^45^. In the myeloid compartment, n=261 osteoclasts expressing *MMP9* and *CTSK* were detected, together with their precursors (Fig2 A, B)^46,47^. Finally, a small number of adipose CXCL12-abundant-reticular (adipo-CAR) cells were captured, expressing *LPL* & *CXCL12* but lacking osteogenic markers other than *ALPL* (Fig. 3G)^48^.

Examining heterogeneity over time, the fraction of cell states in each compartment changed through development (Fig. S3A, C, D). The earlier samples were dominated by chondroprogenitors and early resting chondrocytes, with older samples consisting of more mature cells (Fig. S3C). For the first trimester bone samples, a proximo-distal developmental gradient was observed at each time point, with the femora composed of fewer chondroprogenitors than the tibiae & fibulae (Fig. S3B). For the 8 PCW sample, in which the right femur and tibia were sectioned into proximal and distal segments, a clear gradient of proximo-distal patterning genes was observed (Fig. 3H). Mature chondrocyte marker genes were more strongly expressed in proximal samples, with early mesenchymal progenitor maker genes dominating the distal samples (Fig. 3H). Similarly, differential abundance testing across all samples revealed enrichment of resting chondrocytes in the tibia & fibula, with Prehypertrophic chondrocytes enriched in the femora (Fig. 3I). There were no patterning genes enriched in the right vs left side or vice versa, including the nodal pathway genes, likely due to the relatively advanced stage of sampling.

We also found expression of *MPZ* (a marker of Schwann cell precursors, SCPS) within the cartilage anlage of the hindlimb (Fig. S3F). To explore this further, we performed RNA in situ hybridisation on an adjacent tissue section (PCW6.2) to test the expression of *MPZ* and other classical Schwann cell precursor genes *ERRB3* and *SOX10*. All three genes showed strong staining within the cartilage anlage of the foot plate (Fig. S3E). Analysis of the single cell data also revealed expression of these genes within the chondrocyte lineage, with 8,985 out of 55,030 cells co-expressing *SOX9* along with at least one other SCP marker, and 38 cells expressing *SOX9* with all three SCP markers (Fig. S3G, H). It is unclear whether this reflects that SCPs, as is the case in the axial skeleton, contribute to the chondrocyte lineage, or if these genes simply play an as yet incompletely characterised role in endochondral ossification, however this finding should spur further investigation^49,50^.

### Transcription factor networks through endochondral ossification

We next applied TF network inference to our single cell data using the DoRothEA package for R ^51,52^. This revealed distinct modules of active regulons associated with progression through endochondral ossification (Fig. 3J). For the chondrocyte lineage, a common set of regulons showed increasing activity through chondrogenesis, including *SOX9*, the chief regulator of chondrogenesis, and Kruppel-like factor (*KLF*) 4, which has been shown to regulate matrix-associated genes in chondrocytes ^53^ ^54^. Other chondrogenic regulators that showed steady increases in activity included *STAT3*, *CSRNP1*, *PPARA* and *NKX3.2*^55–58^. Other TFs were specific for certain populations. These included *HAND1,* predicted as being active in chondroprogenitors, which inhibits chondrocyte maturation, and the *MMP13* regulator *NR4A3*, which was most active in prehypertrophic chondrocytes^59,60^. Similarly, *DLX3* & *NFAT5,* which are involved in chondrocyte hypertrophy and osteoblastgenesis, as well as *KLF15*, which promotes *SOX9* activity during *in vitro* chondrogenesis from MSCs, were active in prehypertrophic and hypertrophic chondrocytes ^61–64^. Interestingly, *LBX1*, which is associated with adolescent idiopathic scoliosis, was also predicted to be active in these two subtypes^65^.

The periosteal lineage also exhibited a specific module of regulon activities, including several known regulators of bone development active in all populations such as *HAND2, OSR2, EBF2* and *ZFHX4* (Fig. 3J)^66–69^. *RUNX2* activity was predicted in periosteal, but not perichondrial, cells. *FOSL1,* which causes osteopenia when knocked out in the embryonic phase in mice, was predicted to be active in perichondrium and *MMP9*-expressing periosteal cells^70^.

Perichondrial-derived pre-osteoblasts exhibited distinct TF activities compared with 2nd trimester de facto osteoblasts (Fig. 3J). Whilst the osteoblastic TF *DLX5* was predicted to be active, other regulators also showed activity including *GLI3, POU2F1* & *AHR*^71–74^. Remaining osteoblastic populations exhibited *TFCP2* activity, which is thought to activate *BMP4* in osteoblast maturation ^75^. Interestingly, HEY1-osteoblasts exhibited *ERF* activity, which suppresses *RUNX2*, explaining their lack of *RUNX2* activity and immature transcriptomic profile as compared to early and mature osteoblasts^76^ (Fig. 3G). These latter two cell types had predicted activity in *FOSL2*, which regulates *BGLAP*, as well as the pro-osteogenic TFs *ATF4* and its activator *USF2* ^77,78^. Finally, two members of the FOXO family of transcription factors were found to have differential activities through development. *FOXO1* showed strong activity in early and mature osteoblasts, whereas *FOXO3* was highly active in hypertrophic chondrocytes and early osteoblasts.

### Using embryonic skeletogenesis to benchmark *in vitro* chondrogenesis protocols

We next applied our single cell atlas of *in vivo* bone and joint formation to evaluate a range of publicly available bulk and single-cell RNA sequencing datasets generated from *in vitro* differentiation of chondrocytes from various human sources including embryonic stem cells (hESCs), induced pluripotent stem cells (iPSCs), mesenchymal stem cells (MSCs) and fibroblasts^79–83^(Fig. 4A; Supplementary Table 2). These protocols were sampled at a range of differentiation time points (Fig. 4A; Supplementary Table 2). We also analysed newly generated bulk RNA data from our own protocol that generates chondrocytes through an pluripotent stem cell-mesenchymal cell (iMSC) intermediate (see methods)^84^. To analyse bulk RNA data, we applied the CellSignalAnalysis python package to search for common transcriptional programs between the *in vitro* data and our embryonic single cell reference atlas, and then validating results with analysis of osteochondral marker gene expression (as defined by expression in the fetal limb scRNAseq data) in the bulk data using the edgeR and limma R packages (Fig. 4A, Fig. S4A)^85–87^.

**Figure4.**
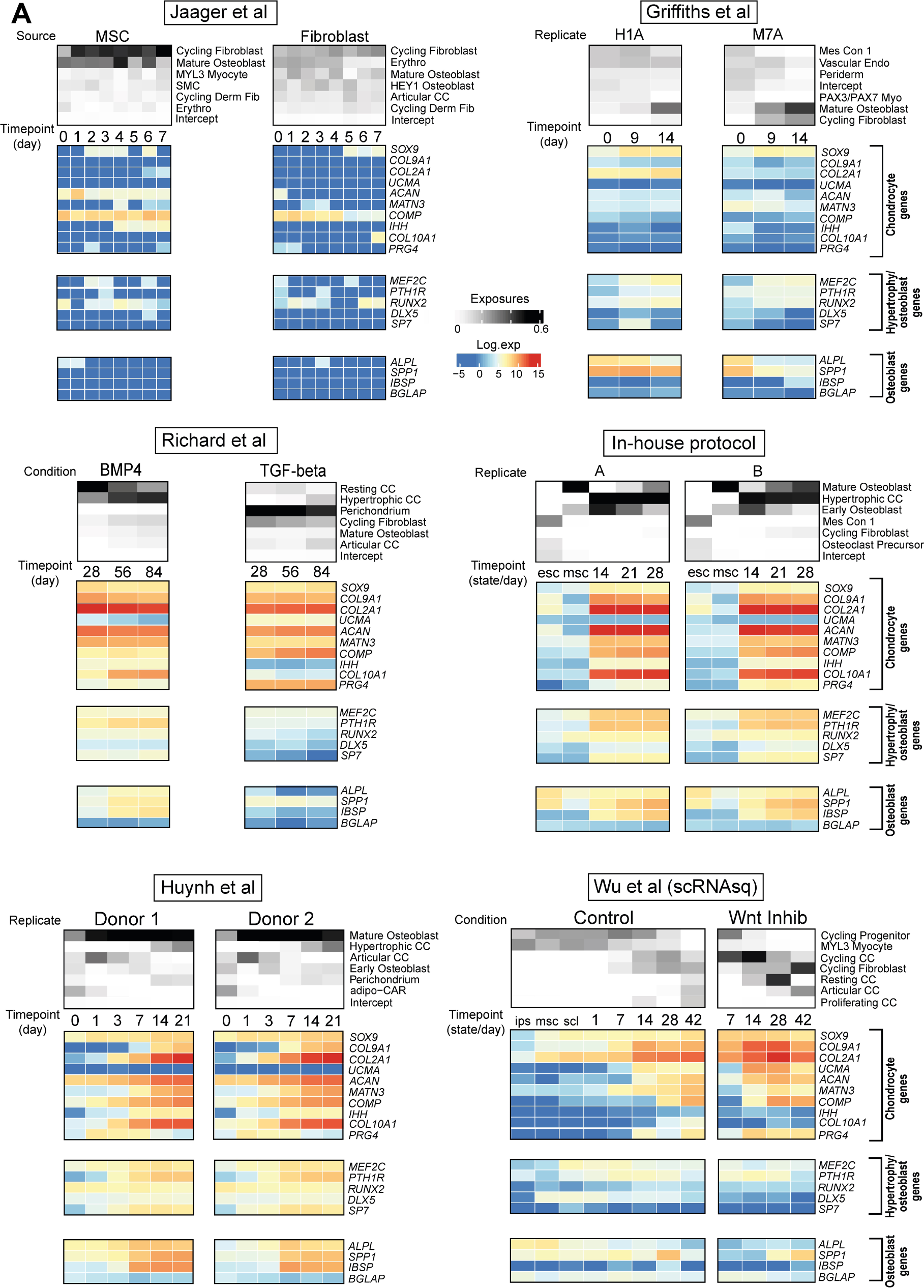
The landscape of *in vitro* chondrogenesis. **(A)** The relative contribution of single cell derived signals from embryonic skeletal tissue in explaining the bulk transcriptomes of samples taken during six *in vitro* chondrogenesis protocols at varying time points, together with marker gene expression. For each protocol, the upper heatmap shows the relative contribution of each embryonic cellular signal to each bulk RNA-seq sample on the y-axis, with sample stage on the x axis. Colour relates to the cell fraction, ranging from zero (white) to 0.6 (black). The lower heatmap shows log-normalised expression of marker genes at each sample stage, ranging from low (blue) to high (red). SMC; Smooth Muscle Cell, Derm Fib; Dermal Fibroblast, Erythro; Erythrocyte; CC; Chondrocyte, Mes Con; Mesenchymal Condensate, Endo; Endothelium, Myo; Myocyte, ESC; Embryonal Stem Cell, MSC; Mesenchymal Stem Cell, IPS; Induced Pluripotent Stem Cell, SCL; Sclerotome

The analysis revealed a heterogeneous landscape of *in vitro* chondrogenesis, with only some protocols producing cells that convincingly shared transcriptional programmes with *in vivo* fetal chondrocytes (Fig. 4A). Two chondrogenic protocols utilised fibroblasts and MSCs respectively, sampling at 24-hour intervals up to day 7 for transcriptomic analysis (Fig. 4A). The MSC-based protocol matched to fibrous and osseous cell types with a low intercept term (that is, the fraction of the bulk transcriptome that cannot be explained by reference cell types), exhibiting only weak expression of *SOX9, COL10A1, ACAN* and *COMP,* but with no other chondrogenic markers being significantly expressed (Fig. 4A). The same protocol applied to a fibroblast cell line produced fewer chondrocyte genes, with consequent non-specific cell-type matching (Fig. 4A). Similarly, a protocol from Griffiths et al designed to generate chondroprogenitors that sampled at the MSC stage (day 0), day 9 and day 14 of differentiation matched to fibro-osseous tissue and exhibited diffuse, weak chondrogenic gene expression, with osteoblast genes *SPP1 & ALPL* upregulated (Fig. 4A). Four further protocols did exhibit strong chondrogenic gene expression patterns, with consequent matching to embryonic chondrocyte populations (Fig. 4A). An 84-day BMP4-supplemented protocol by Richard et al. sampled at days 28, 56 and 84 matched to hypertrophic chondrocytes and exhibited strong *COL10A1* and *IHH* expression at all time points, whilst the same group’s TGF-beta-supplemented protocol exhibited a stronger articular signal, with corresponding *PRG4* expression, without *COL10A1* or *IHH* (Fig. 4A). Similarly, our 28-day in-house hESC protocol, which was sampled at the hESC & MSC stages, followed by days 14, 21 & 28 of differentiation, showed a clear shift to a hypertrophic chondrocyte gene module (including *COL10A1* and *IHH*) at day 14 onwards (Fig. 4A). The same was true of an iPSC protocol from Huynh et al., which was sampled more densely at early time points (days 0, 1, 3, 7, 14, 21 of chondrogenic differentiation) and showed a steady increase of the same transcripts (Fig. 4A). Both protocols also showed strikingly similar expression of transcripts relating to hypertrophy and osteoblastgenesis (*MEF2C, PTH1R, RUNX2, DLX5 & SP7*) as well as specific osteoblastic transcripts *ALPL, SPP1* and *IBSP* (Fig. 4A). Consequently both showed strong matches to both hypertrophic chondrocytes and osteoblasts (Fig. 4A). Both in-house and Huynh et al’s protocols also exhibited *PRG4* expression, though at lower levels than those produced by Richard et al. and with no cell signal matching to articular chondrocytes (Fig. 4A). Finally, a single cell study by Wu et al. of an iPSC protocol showed similar gradual upregulation of chondrogenic genes over time, however in this case *UCMA* and *PRG4*, rather than *COLA10A1*/*IHH* were expressed, with consequent matching to proliferating, resting and articular chondrocytes (Fig. 4A). *UCMA* and *PRG4* expression together with resting and articular chondrocyte signals were increased with the addition of the small molecule Wnt inhibitor WNT-C59 (Fig. 4A). Analysis of predicted transcription factor (TF) network activity showed that protocols predicted to match to embryonic osteochondral populations exhibited activity in many of the same TFs as their *in vivo* counterparts (FigS5).

### Chondrocyte Differentiation Dynamics *in vivo* and *in vitro*

To compare differentiation dynamics *in vivo* and *in vitro* at single-cell resolution and enable assessment of alterations to culture conditions throughout differentiation, we performed 10X chromium-based single-nuclei RNA sequencing (snRNAseq) on newly generated *in vitro* cells from our in house protocol (Fig. 5A; see methods). We harvested cells from independent replicate experiments at multiple differentiation timepoints from the MSC stage onwards. Following pre-processing and quality control (see methods), we captured 153,524 single nuclei (Fig. S6A for QC).

**Figure5.**
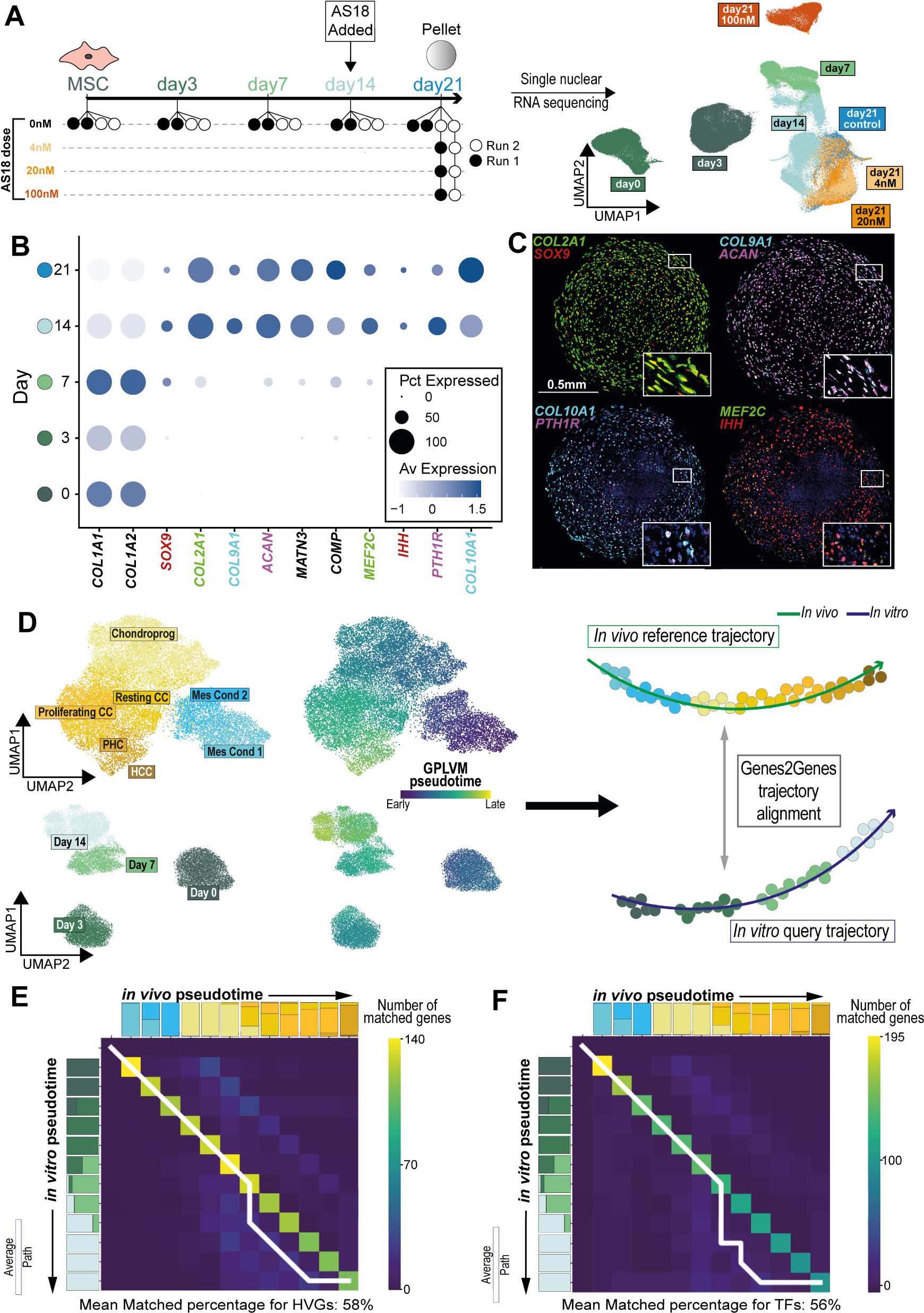
Chondrocyte differentiation *in vitro* and *in vivo*. **(A)** left panel-Experimental Overview. Each circle corresponds to a separate sequencing batch. Right panel-Uniform Manifold Approximation & Projection (UMAP) plot of 153,524 single nuclei from *in vitro* chondrocytes. **(B)** Dotplot visualisation showing scaled marker gene expression for cells depicted in (A). Dot size corresponds to the fraction of cells with non-zero expression. Label colours on the x-axis correspond to staining colours in (C). Dots are coloured according to gene expression level, ranging from not detected (white) to the highest detected levels (blue), according to the adjacent colour key. **(C)** RNA-In Situ Hybridisation of sections of *in vitro* hPSC-chondrocyte pellets taken at day 21, for *COL2A1, SOX9, COL9A1, ACAN, COL10A1, PTH1R, MEF2C* and *IHH*. Inset high magnification boxes represent the regions in the smaller boxes on the lower magnification image. **(D)** left panel-Uniform Manifold Approximation & Projection (UMAP) plots for *in vivo* (upper plots) and *in vitro* (lower plots) cells utilised for trajectory alignment, together with GPLVM their pseudotime values, as indicated by the adjacent colour scale. **(E)** Pairwise time point matrix between *in vivo* and *in vitro* pseudotime for highly variable genes. The colour represents the number of genes showing match or warp for the given pair of an *in vitro* time point and an *in vivo* time point. The white line represents the main average alignment path. HVG; highly variable genes **(F)** Pairwise time point matrix between *in vivo* and *in vitro* pseudotime for transcription factors. The colour represents the number of genes showing match or warp for the given pair of an *in vitro* time point and an *in vivo* time point. The white line represents the main average alignment path. TFs; transcription factors

Cells at day 0 & 3 formed distinct clusters, with days 7, 14 & 21 showing a progression through chondrogenesis (Fig. 5A, B). At day 7, subtle differences in the differentiation state of cells were observable, with less differentiated cells (Progenitor 1 & 2) more strongly expressing *COL1A1* & *COL3A1,* and more differentiated cells (Chondro 1, 2 & 3) expressing *COL2A1* (Fig. S6D). One progenitor cluster (Progenitor 1) also strongly expressed *PLCG2*, in keeping with a previous study showing downregulation of this gene in MSCs as they begin chondrogenesis (Fig. S6D)^88^. A cluster of mature cells (Chondro 1) expressed β-catenin (*CTNNB1*), a known regulator of chondrocyte maturation, with a second expressing *PENK* (Fig. S6D)^89^. These genes were maximally expressed in prehypertrophic and proliferating fetal chondrocytes respectively (Fig. S6E). At day 14, most cells were expressing *COL10A1*, supporting the findings of the bulk-RNA analysis (Fig. 5B). However, the expression varied between different runs, with run 2 being more differentiated than run 1 at this time-point (Fig. S6B, C). By day 21 this batch difference was no longer evident (Fig. S6B, C). One cluster of day 21 cells expressed higher *COL10A1, COMP, IBSP* and *FMOD*, indicative of further maturation at this time point (Fig. S6B, C; red dashed line; Supplementary Table 3 for DEGs). RNA FISH at day 21 confirmed expression of several markers of chondrogenesis and chondrocyte hypertrophy (Fig. 5C). Whilst pan-chondrocyte markers (*SOX9, COL2A1, COL9A1, ACAN)* were expressed throughout the pellets, expression of hypertrophic markers (*COL10A1, PTH1R, IHH, MEF2C*) was most intense in the periphery (Fig. 5C)

To compare *in vivo* and *in vitro* differentiation dynamics, we performed trajectory alignment using a dynamic programming algorithm implemented by Genes2Genes^90^ python package, treating *in vivo* chondrocyte lineage data (mesenchymal condensate through to hypertrophic chondrocyte) as the reference, and *in vitro* cells from MSC to day 21 as the query (Fig. 5D; see Methods). This enabled each gene to be aligned independently, where we obtain optimal sequential mapping between the *in vitro-in vivo* pseudotime points for each gene. This enabled us to categorise genes at each point in pseudotime with how closely they align in each trajectory as a match or a mismatch (see methods). This approach revealed moderate overall alignment of the two trajectories, whilst highlighting large numbers of transcripts (both TFs and non-TFs) which were aberrantly expressed *in vitro* (Fig. 5E & F; Supplementary Tables 4 & 5). Across all highly variable genes, the mean percentage of trajectory time points where expression *in vivo* and *in vitro* matched was 58% (Fig 5E). This remained similar for TFs (Fig. 5F). Indeed, several key chondrogenic transcription factors exhibited matched across the two trajectories, including *SOX6, SOX9*, *KLF2*, KLF*4*, *FOXP1*, *STAT3*, *NKX3-2* and the mesenchymal TF *HMGA2* (Fig 6A; Supplementary Table 5). This trend was accompanied by high proportions of matches in many classical non-TF chondrocyte transcripts, including *COL2A1, ACAN, MATN3,COL9A2, COL9A3* and *IHH* although a small number still exhibited poor matching, including *EPYC, COL9A1*, *MATN1* and *MATN4* (Fig. 6B; Supplementary Table 4). These analyses also revealed how the expression of large numbers of off-target transcripts that were aberrantly expressed *in vitro*, including the smooth muscle gene actin alpha-2 (*ACTA2*), the cilia-related gene *CFAP298*, limb mesenchymal regulator *PITX1* (absent *in vitro*), the perichondrial gene *THBS2* and the osteoblastic genes *COL1A1*, *ALPL*, *SPP1, LRP4, SP7* and *FOXO1* (Fig. 6C; Supplementary Tables 4 & 5). This likely explains why analysis of the bulk transcriptome with CellSignalAnalysis revealed signals from non-chondrocyte cell types, particularly osteoblasts (Fig. 4A).

**Figure 6.**
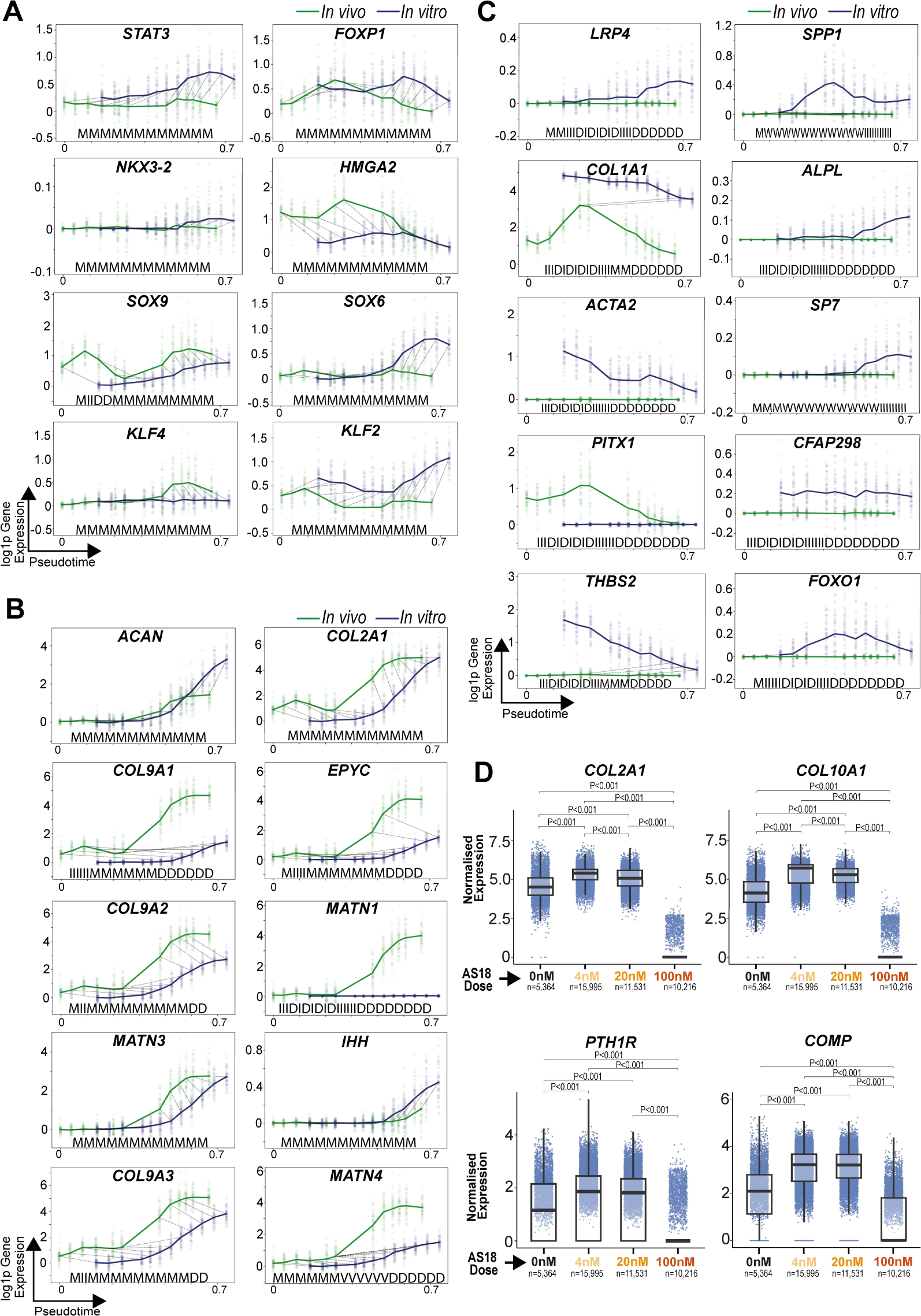
Gene expression dynamics during chondrogenesis *in vitro* and *in vivo*. **(A)** Plots showing interpolated log-normalised expression (y-axis) against pseudotime (*x*-axis) for selected transcription factors for *in vitro* (blue) and *in vivo* (green). The bold lines represent mean expression trends, while the faded data points are 50 random samples from the estimated expression distribution at each time point. The black dashed lines represent matches and warps between time points. The five-state alignment string for each gene is shown below the expression plots. M; Match, W; many-to-one match, V; one-to-many match, I; insertion, D; deletion. **(B)** The same plots as (A), showing the expression of classical chondrocyte genes. **(C)** The same plots as (A) & (B), showing the expression of genes aberrantly expressed *in vitro*. **(D)** Box plots showing the expression of chondrocyte genes at different AS18 concentrations. The box contains the 25th to 75th percentiles of the data, with the central line denoting the median value. The upper whisker extends from the median to the largest value, no further than the 1.5 * inter-quartile range (IQR). The lower whisker extends from the median to the smallest value, at most 1.5 * IQR. Wilcoxon rank-sum test, two-sided, with adjustment for multiple comparisons.

Examining the snRNAseq data, we found that the aforementioned off-target transcripts were diffusely expressed throughout the manifold, including in cells that were positive for canonical chondrocyte genes (Fig S6F). This contrasts with a previous study which found distinct clusters of off-target cells that pursued different lineages, with little expression of chondrocyte genes^80^. In summary, our differentiation protocol appears to produce chondrocytes that aberrantly express some off-target transcripts, rather than a mixture of chondrocytes and undesirable cells, such as neural crest cells, melanocytes or neuronal-like cells.

To reduce off-target (osteoblastic) differentiation, we next searched for TFs active in both embryonic osteoblasts and *in vitro* cells for which well-characterised inhibitors are already available. One TF identified was *FOXO1*, which is thought to promote *RUNX2* activity in mesenchymal stem cells ^91^. An inhibitor of FOXO1, AS1842856 (“AS18”), is available, making it an appealing target for manipulation. AS18 exhibits specificity for FOXO1 at low concentrations (IC50 = 33nM), but inhibits FOXO3&4 family members at higher concentrations (IC50 > 1µM). We hypothesised that FOXO1 blockade in less mature cells (day 14) may prevent the off-target osteoblastic differentiation thereby increasing chondrogenic gene expression and improving chondrocyte yield (Fig. 5A).

We first performed an AS18 dose-response using validated BMP and TGFꞵ signalling reporter cell lines to identify the concentrations that may regulate chondrogenesis^92–95^. This showed increased BMP signalling and decreased TGFꞵ signalling upon AS18 addition, up to a threshold (Fig. S7A). Based on the reporter cell line results, we added AS18 at several concentrations (4 nM, 20 nM or 100 nM) at day 14 of the protocol. The resulting manifold projection showed control, 4 nM and 20 nM populations clustering together, with 100 nM forming a distinct cluster (Fig. S7B). Differential expression testing at day 21 revealed that cells cultured with low concentrations of AS18 (4 nM, 20 nM) exhibited an increased expression of chondrocyte genes, such as *COL2A1, COL9A1, MATN3,* and *COMP*, together with markers of hypertrophy, in both differentiation runs when compared to controls (Fig. 6D; Fig. S7C; Supplementary Table 6 for DEG lists). At 4 nM and 20 nM concentration, the latter included *COL10A1, PTH1R, FGFBP2, SPP1,* and *FMOD* (Fig. 6D; Fig. S7C; Supplementary Table 6). *ACAN* and *SOX9* were upregulated at 4 nM concentration only (Fig. S7C; Supplementary Table 6). The BMP response gene *ID1* was significantly upregulated at 20 nM and 100 nM, in keeping with the dose-response observed in the reporter cell line (Fig. S7C). Some osteoblastic genes were also down regulated including the *RUNX2* co-activator *DDX5* together with the pro-osteogenic lncRNAs *KCNQ1OT1*, and *SNHG14* (Fig. S7D)^96–98^. Interestingly, *COL27A1* was also downregulated; a gene implicated in the calcification of the cartilage anlage^99^. Cells cultured with 100 nM AS18 concentration showed minimal chondrogenic gene expression, suggesting our cells exhibited increased sensitivity to AS18 when compared to the reporter cell lines; a finding likely explained by their greater expression of FOXO family members compared to these reporters (Fig. 6D; Fig. S7A & C). Although 100nM AS18-treated cells expressed few chondrogenic genes, this concentration did not appear to be toxic; of the 87 KEGG-pathway genes for apoptosis, only 4 (*IKBKB, NFKBIA, BAX, & EXOG*) were differentially expressed in these cells compared to controls (Supplementary Table 6; see methods).

## Discussion

In this study, we generated a detailed embryonic reference atlas to evaluate the current landscape of *in vitro* chondrogenesis. Through analysis of engineered cartilage against our reference, we quantified both on- and off-target differentiation *in vitro* at the single-cell level, giving new insights into how culture conditions may stimulate transcription of some classical chondrogenic genes whilst failing to induce the global expression patterns in cells required to faithfully recapitulate *in vivo* development. This detailed interrogation of engineered tissue may serve as a framework for future work both in cartilage and other tissues. In addition, we identified pathways that could be manipulated to push *in vitro* cells into a desired state. Through inhibition of *FOXO1*, a molecule implicated in osteogenesis, we increased chondrocyte gene expression *in vitro*. This serves as a proof of principle for utilising data from human development to further tissue engineering in the musculoskeletal system, building on the work in other fields ^4,5^.

We envisage future work using these approaches will enable us to fine-tune protocols through the targeting of a range of molecules implicated in embryonic chondrogenesis, minimising off-target transcription and more faithfully recapitulating *in vivo* cell states. Whilst *FOXO1* was an attractive target due to its high activity in the osteoblastic lineage and the existence of a readily available inhibitor, it was predicted to be active at low levels activity in mature chondrocytes. This perhaps explains why the increase in some hypertrophic chondrocyte transcripts was most pronounced at the 4 nM AS18 dose; higher concentrations may have inhibited chondrocyte hypertrophy. Other TFs that currently lack validated inhibitors, but exhibit greater cell-type specificity, would be desirable targets for future studies.

We identified only a limited number of chondrogenesis protocols that have been investigated at the bulk or single-cell transcriptome level which had publicly available data, with many studies analysing only a restricted number of transcripts. This in itself is a major barrier to high fidelity tissue engineering. Furthermore, a wide variety of culture conditions, sampling times and cell sources have been used. Whilst some protocols had matching samples at particular time (in particular, day 7 and 14 were common), no two had identical sampling strategies for sequencing. This limited our ability to survey the landscape of *in vitro* chondrogenesis using our methods, and generally prevents any reliable comparison of particular culture conditions or cell sources.

Although single cell sequencing has proven to be an invaluable tool in forming tissue ‘atlases’, it has several limitations. In the context of human development, the cell types captured and sequenced are understandably constrained by the logistics of termination of pregnancy. It is not feasible, for example, to study the nascent human limb bud, which may prove valuable for comparison to early *in vitro* cells, which tended to give less defined signal matching in comparison. Similarly, whilst we captured osteoblasts expressing makers of maturity, and labelled them as “mature”, the sampling window for public data used in this study did not extend beyond PCW17, thus further cellular maturation was not captured. A further technical limitation arises when investigating matrix-rich or mineralised tissues. In this setting, the breadth of cell capture is constrained by the effectiveness of tissue dissociation into single cell suspension, with cells embedded in a dense matrix less likely to be liberated. This perhaps explains why comparatively few hypertrophic chondrocytes were sequenced (n=61) compared to resting (n=6,852), despite them being histologically abundant during the sampling window. It is indeed possible that the most mature hypertrophic chondrocytes were not captured. The resulting gap in the reference atlas would then increase the portion of the bulk transcriptome not attributable to fetal cell types. It is conceivable that an elusive subset of fetal hypertrophic chondrocytes expressing greater levels of transcripts that have biological overlap with osteoblasts, (such as RUNX2/3 or DLX5) or indeed cells undergoing transdifferentiation to the osteoblastic lineage, could match to bulk transcriptomes and lower the portion predicted to be attributable to fetal osteoblasts^100^. Indeed, the protocols with available bulk sequencing analysed in this study which have been analysed histologically or by electron microscopy do show chondrocyte characteristics^79,80,83^. This fact highlights the broader question of the importance of off-target transcripts in determining the characteristics of the tissue produced. It may be that, for example, aberrant expression of *ALPL* has minimal functional consequence on the tissue produced. In other words, perhaps a degree of off-target transcription can be accommodated in the pursuit of high-fidelity cartilage engineering.

A key issue for engineering cartilage is defining the desired product of a chondrogenesis protocol. In this study, we showed that ESCs and MSCs will form hypertrophic or resting chondrocyte-like cells under specific conditions, with realistic cartilage matrix transcript expression, albeit with some off-target transcription. However, whilst this progression from progenitor to hypertrophic chondrocyte is invaluable for modelling developmental skeletal diseases, a different phenotype, such as PRG4-expressing articular chondrocytes, may prove more valuable in the study of degenerative joint diseases such as osteoarthritis. For example, the protocol employed by Richard et al. demonstrated that creating culture conditions with different dominant signalling pathways can produce chondrocytes with either an articular or endochondral ossification phenotype^83^. Investigating this approach with high resolution techniques, as employed by us here, could yield further insights into the control of chondrogenesis *in vitro*, enabling modelling of a diverse set of conditions and bringing the prospect of engineered articular cartilage as a therapy closer to the clinic.

One limitation of the *in vitro* component of this study is the inter-experiment variability observed during chondrogenesis. Although our protocols reliably produced hypertrophic chondrocytes, snRNAseq analysis revealed that the rate of differentiation varied between the two runs after the day 7 timepoint (although converging again at day 21), with run 2 exhibiting greater expression of hypertrophic transcripts. This observation was not limited to our own protocol, and is a challenge for tissue engineering in general^101,102^. Furthermore, the duration of chondrogenic differentiation protocols, even from the MSC stage, is significant. This makes high frequency sampling and sequencing a practical difficulty, forcing researchers to choose particular time points based on previous experimental work. Here, based on initial findings from bulk RNA cell signal analysis, we chose to sample at days 0, 3, 7, 14 and 21. However the change in transcriptome from days 7 to 14 was profound; progressing from very weak *SOX9* and *COL2A1* expression to full hypertrophy with abundant *COL10A1* expression. It is not clear if more frequent sampling between these timepoints would reveal a steady progression through resting, proliferating and prehypertrophic states, or if *in vitro* differentiation in this case is more direct from progenitor to hypertrophic cell. Future densely sampled studies would require vast numbers of parallel differentiation experiments to be established, with significant resources required to perform large-scale sequencing at tight time intervals. Similarly, although we employed a reporter cell line to optimise AS18 concentrations, our ESC-derived cells appeared to be much more sensitive than expected from this preliminary work, with *COL2A1* only increasing at low concentrations. We also did not observe decreased *TGF-*ꞵ expression in our cell lines in response to the inhibitor. Although we have shown that there is a switch in the prevalence of gene expression after AS18 application, with some increase in hypertrophic genes, we have not yet shown that this is translated into a greater abundance of cartilage proteins, proteoglycans and GAGs. For this we would need a detailed biochemical analysis which will be the subject of future work. Indeed, whilst we have been able to demonstrate similarities and differences between embryonic and *in vitro* chondrocytes, these analyses were performed at the transcriptome level only, and assessment at the protein level and of the mechanical properties of the product would shed further light on its similarity to *in vivo* tissue.

Musculoskeletal disease is the leading cause of disability worldwide and includes younger people who would normally be working all of which represents a major economic burden for society^6,7,103,104,105^. Understanding the development and homeostasis of these tissues using the methods applied in this study will be a crucial part of both understanding disease pathophysiology and developing new, cell-based treatments for early intervention^106^.

**Figure S1.**
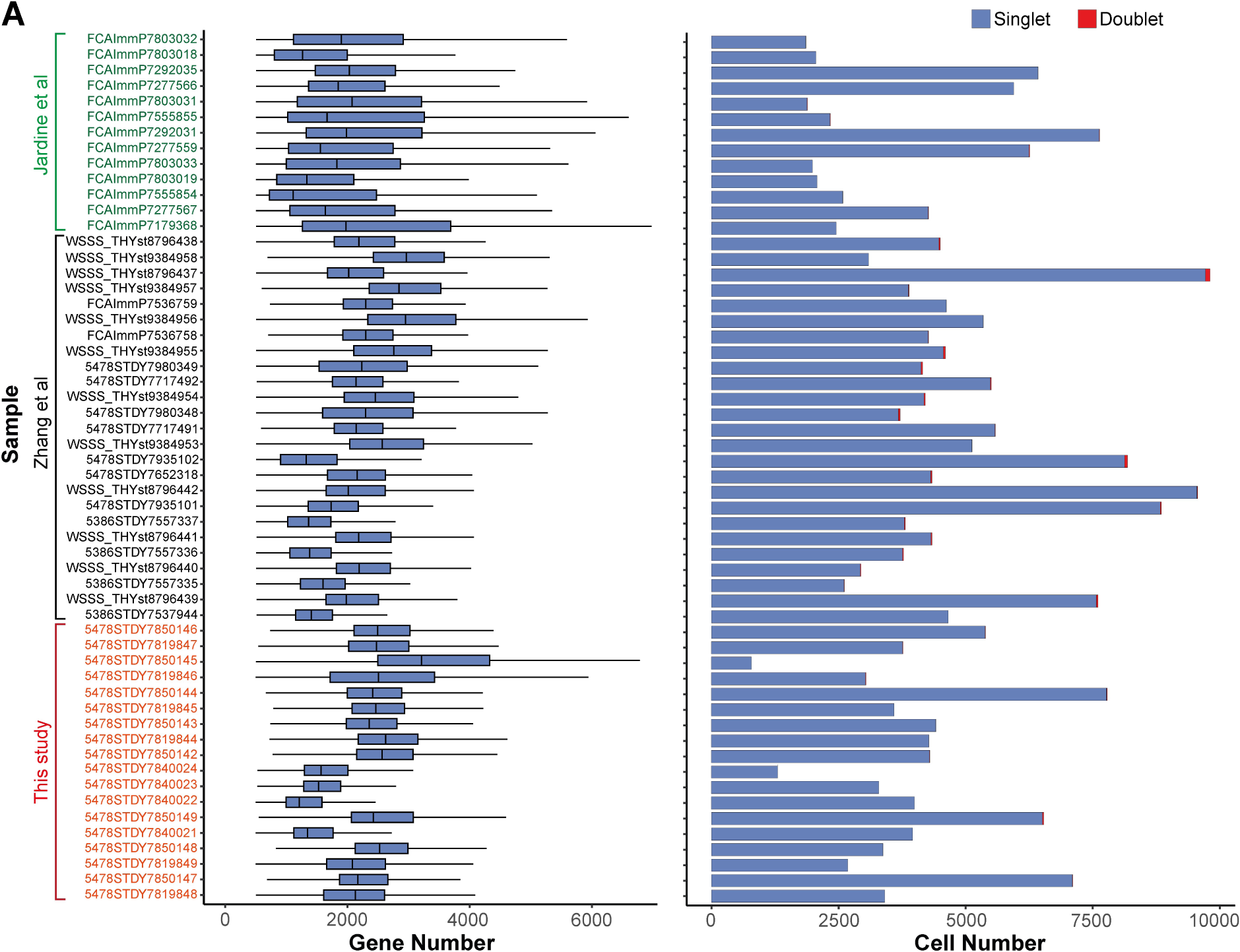
Quality control of *in vivo* data. **(A)** Left panel - Box plot showing per cell gene number for each library. The box contains the 25th to 75th percentiles of the data, with the central line denoting the median value. The upper whisker extends from the median to the largest value, no further than the 1.5 * inter-quartile range (IQR). The lower whisker extends from the median to the smallest value, at most 1.5 * IQR. Right panel - bar plot showing cell number for each library. Red fill indicated cells called as doublets and excluded from further analysis.

**Figure S2.**
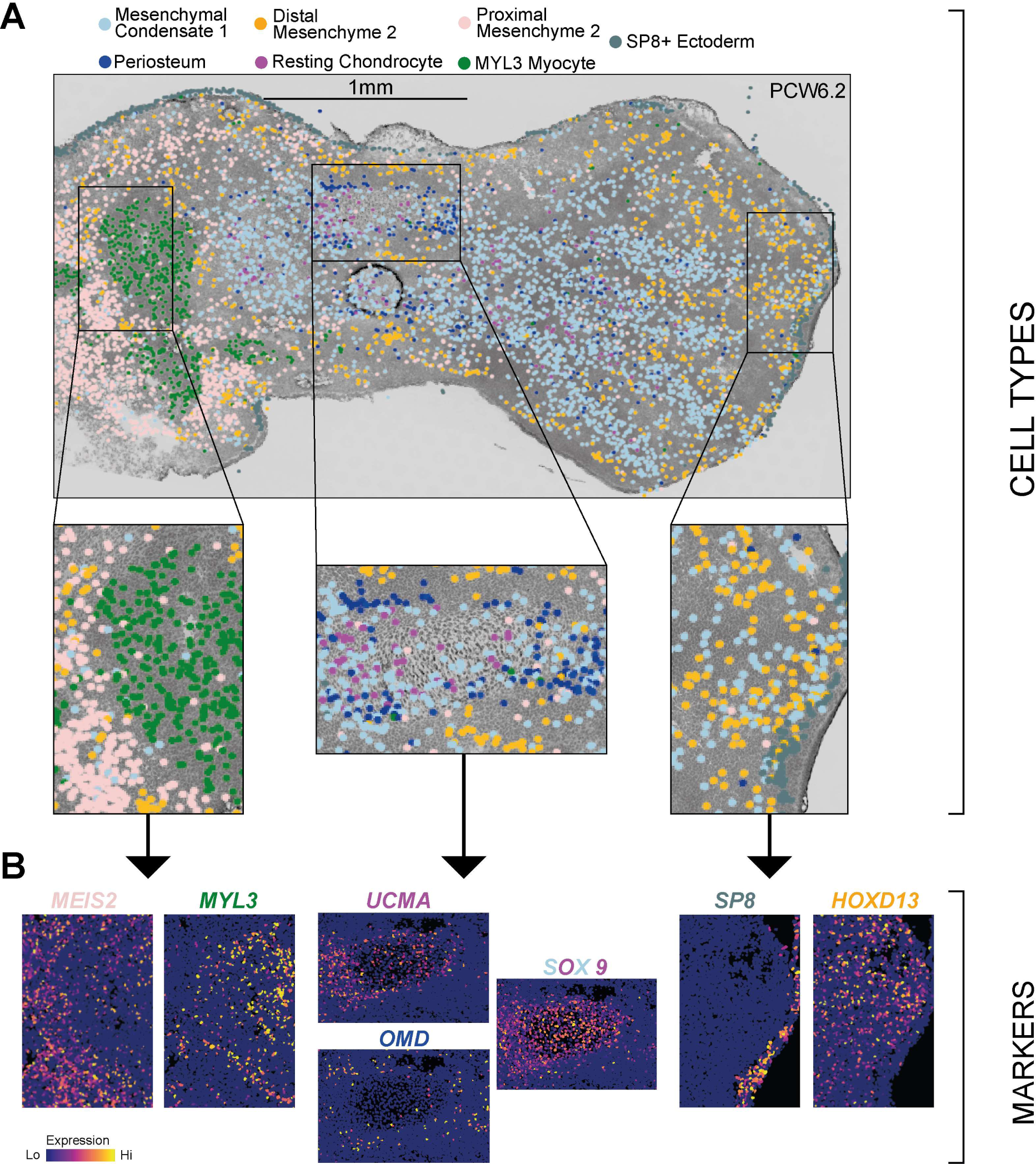
Imputed locations of cells in anatomical space. **(A)** Inferred location of single cells in the PCW6.2 hind limb using k-nearest neighbour prediction on in-situ sequencing data. Coloured points represent predicted cell type location as shown in the legend. Boxed regions represent areas shown at greater magnification in the lower panel. **(B)** Marker gene expression from in-situ sequencing for cell types depicted in (B). Font colour corresponds to cell type marked by that gene. Cells are colour-coded according to gene expression level, ranging from not detected (blue) to the highest detected levels (yellow), according to the adjacent colour key.

**Figure S3.**
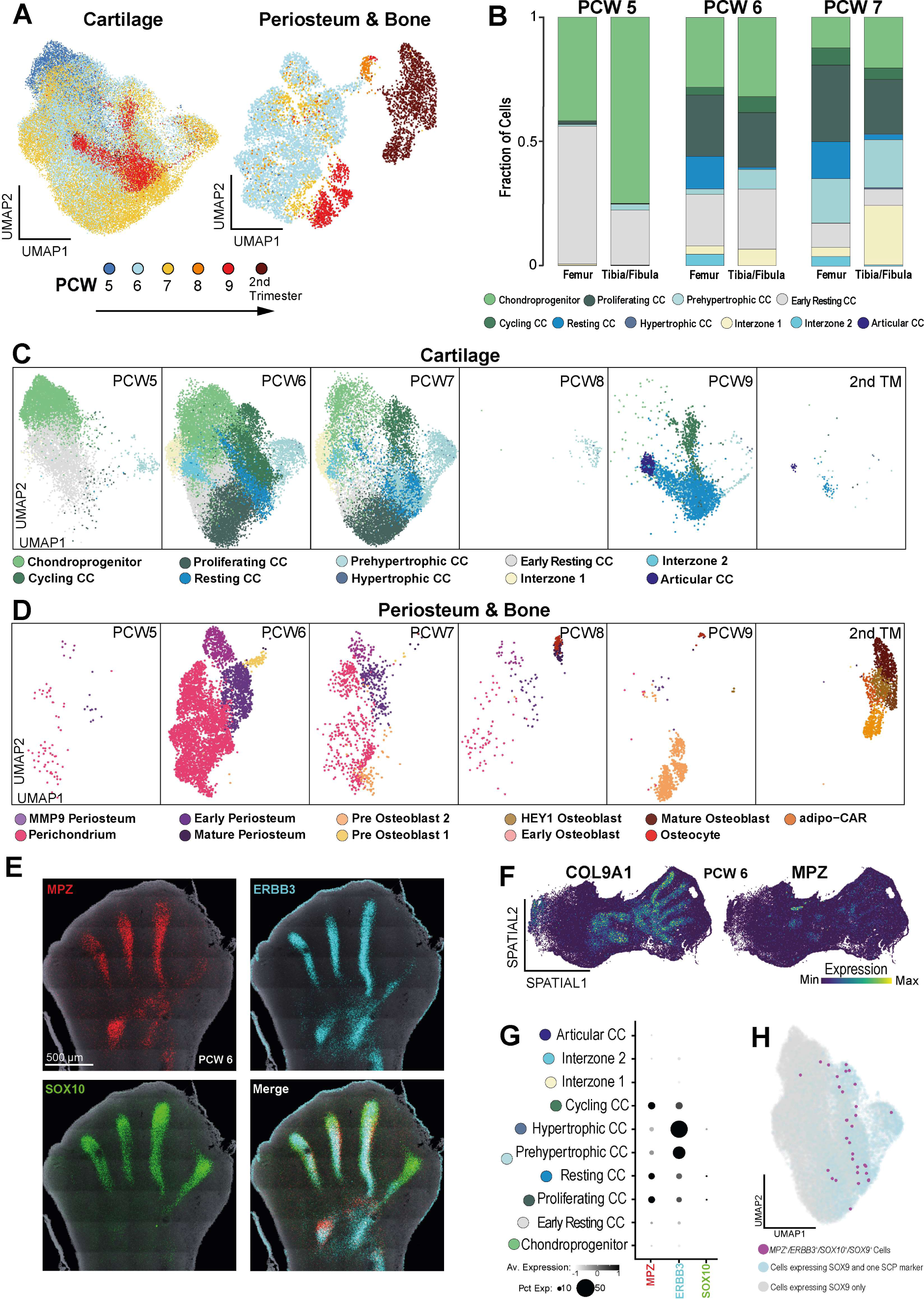
Analysis of the osteochondral compartment. **(A)** Uniform Manifold Approximation & Projection (UMAP) plots of cartilage (left panel) and periosteum & bone (right panel) coloured by sample stage, as indicated in the key. PCW; post-conception week. **(B)** Stacked bar charts showing the fraction of each cell type at each sample stage in the femora and the tibiae/fibulae. CC; chondrocyte, PCW; post-conception week. **(C)** Uniform Manifold Approximation & Projection (UMAP) plots of cartilage cells, split by sample stage. CC; chondrocyte. PCW; post-conception week. **(D)** Uniform Manifold Approximation & Projection (UMAP) plots of periosteum and bone cells, split by sample stage. PCW; post-conception week. **(E)** RNA In Situ Hybridisation of PCW6 foot plate in coronal section showing Schwann cell precursor marker gene expression, *Mpz, ERBB3* and *SOX10*.PCW; post-conception week. **(F)** Spatial heatmap of in-situ sequencing data showing expression of *COL9A1* and *MPZ* in the PCW6 lower limb. Colour corresponds to expression level, as indicated by the key. PCW; post-conception week. **(G)** Dotplot visualisation showing scaled gene expression for *in vivo* chondrocytes. Dot size corresponds to the fraction of cells with non-zero expression. Dots are coloured according to gene expression level, ranging from not detected (white) to the highest detected levels (black), according to the adjacent colour key. **(H)** Uniform Manifold Approximation & Projection (UMAP) plot of *in vivo* chondrocytes showing cells that co-express of *SOX9* with one Schwann cell precursor marker (blue); three schwann cell precursor markers (magenta) or no schwann cell precursor markers (grey).

**Figure S4.**
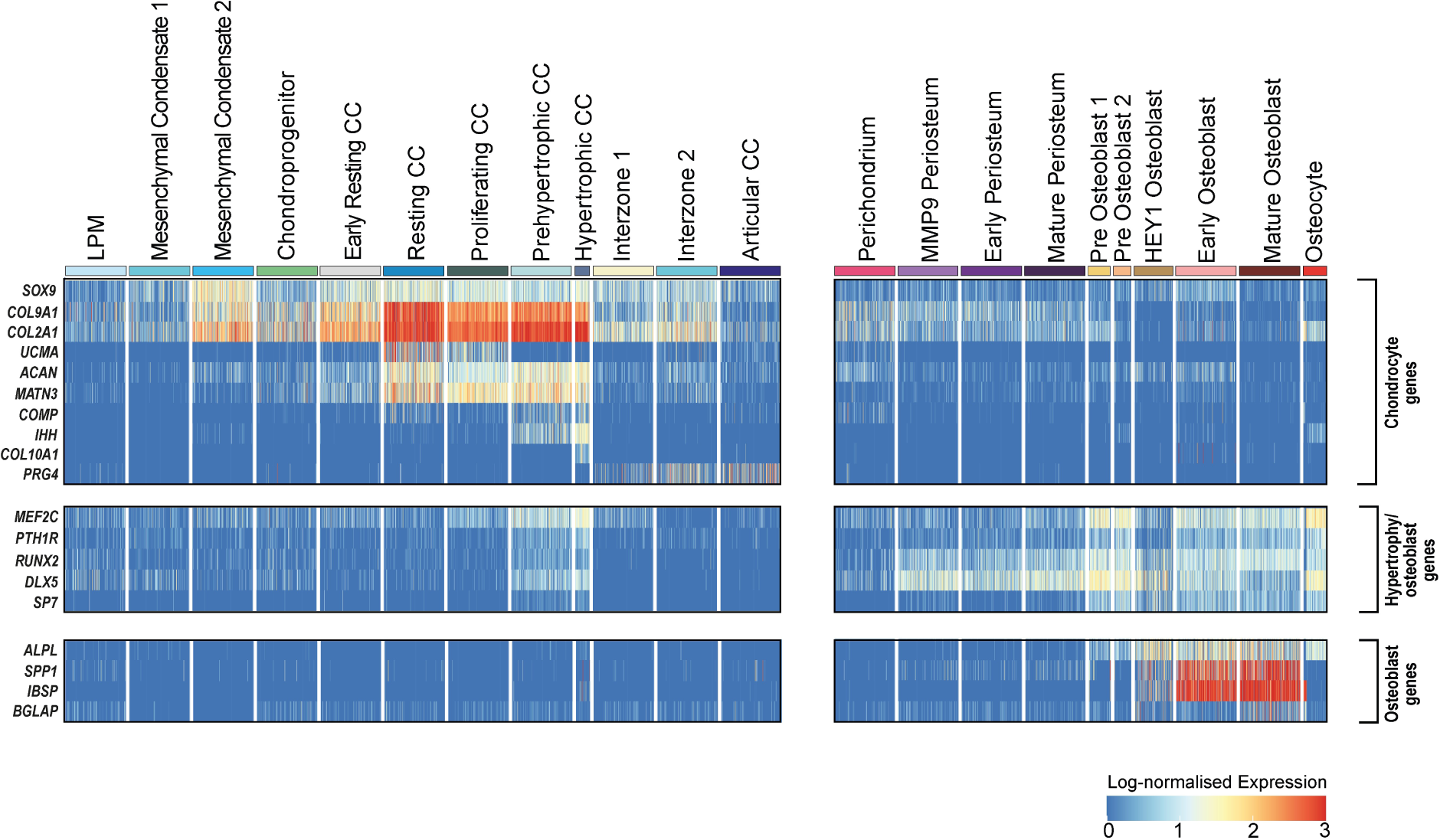
Gene expression across single cells of the fetal limb. **(A)** Heatmap showing log-normalised expression of marker genes used in Figure 4 in fetal osteochondral populations, as indicated by the colour scale. Each column represents a single cell, with cells of the same type grouped into larger columns, as indicated by the colour bars and labels above each.

**Figure S5.**
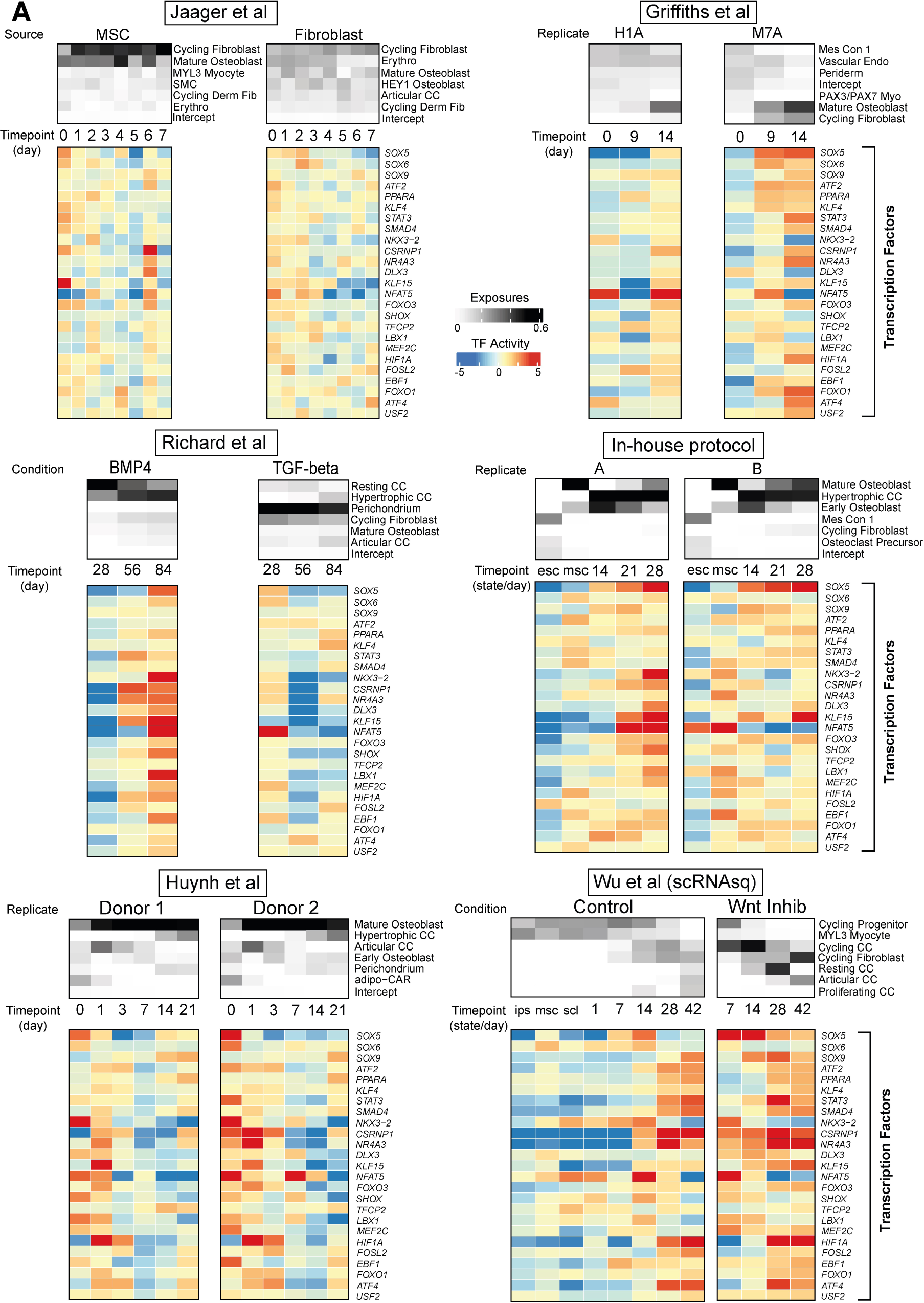
Predicted transcription factor activities in *in vitro* chondrogenesis protocols. **(A)** The relative contribution of single cell derived signals from embryonic skeletal tissue in explaining the bulk transcriptomes of samples taken during six *in vitro* chondrogenesis protocols at varying time points, together with predicted transcription factor activities. For each protocol, the upper heatmap shows the relative contribution of each embryonic cellular signal to each bulk RNA-seq sample on the y-axis, with sample stage on the x axis. Colour relates to the cell fraction, ranging from zero (white) to 0.6 (black). The lower heatmap shows predicted transcription factor activities at each sample stage, ranging from low (blue) to high (red). SMC; Smooth Muscle Cell, Derm Fib; Dermal Fibroblast, Erythro; Erythrocyte; CC; Chondrocyte, Mes Con; Mesenchymal Condensate, Endo; Endothelium, Myo; Myocyte, ESC; Embryonal Stem Cell, MSC; Mesenchymal Stem Cell, IPS; Induced Pluripotent Stem Cell, SCL; Sclerotome

**Figure S6.**
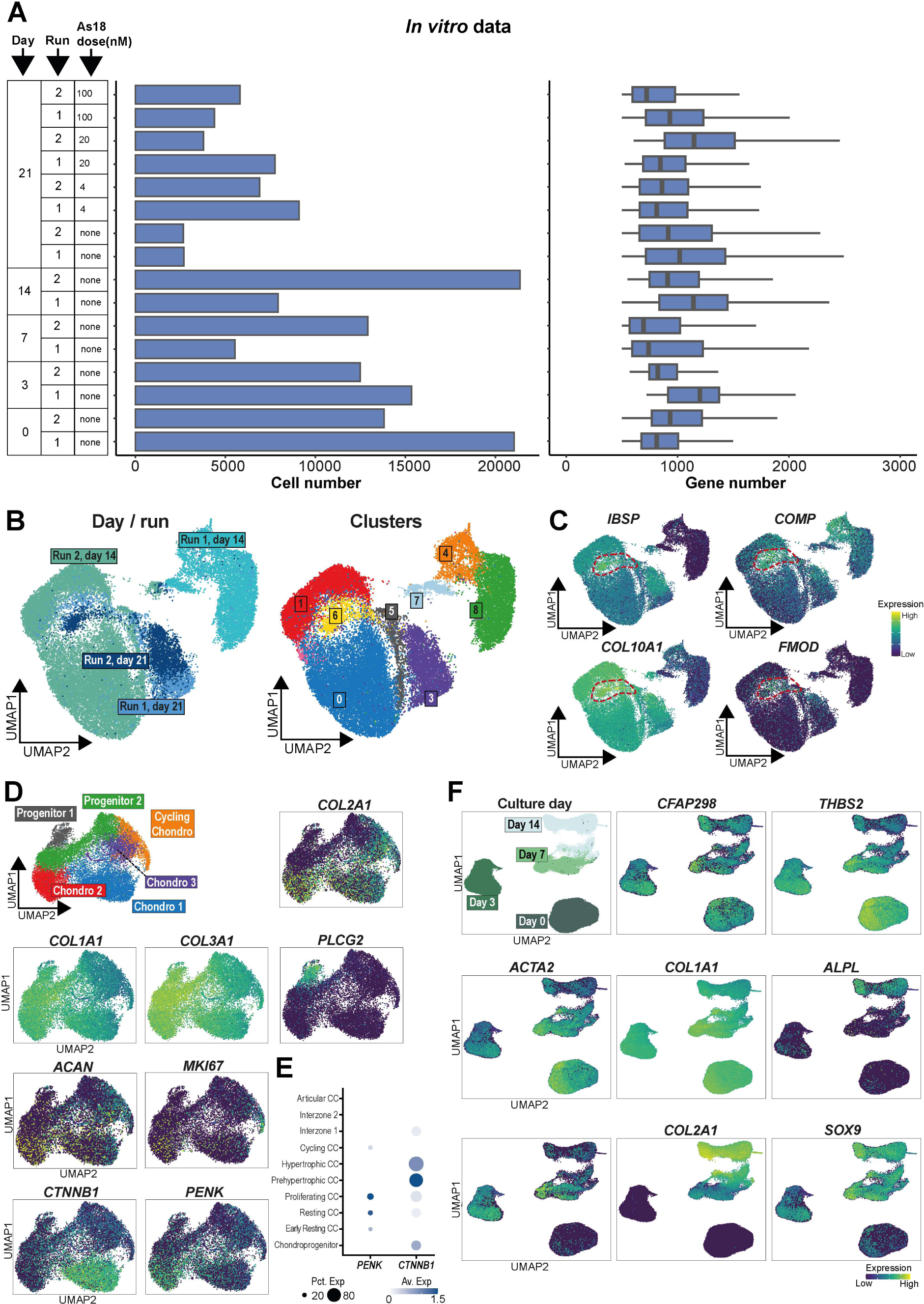
Quality control and gene expression trends for single-nuclear sequencing of *in vitro* chondrogenesis. **(A)** Left panel - bar plot showing cell number for each library. Red fill indicated cells called as doublets and excluded from further analysis. Right panel - Box plot showing per cell gene number for each library. The box contains the 25th to 75th percentiles of the data, with the central line denoting the median value. The upper whisker extends from the median to the largest value, no further than the 1.5 * inter-quartile range (IQR). The lower whisker extends from the median to the smallest value, at most 1.5 * IQR. **(B)** Uniform Manifold Approximation & Projection (UMAP) plots of day 14 and day 21 *in vitro* single nuclei. Left panel - plot showing batch differences, right panel - clustering of cells. **(C)** Uniform Manifold Approximation & Projection (UMAP) plots showing gene expression in day 14 & 21 *in vitro* single nuclei. Red dotted line denotes cluster with stronger expression of hypertrophic transcripts. Dots coloured by expression as shown by the colour bar **(D)**Uniform Manifold Approximation & Projection (UMAP) plots of day 7 *in vitro* single nuclei. Upper left panel - annotated clusters. Remaining panels show gene expression, with dots coloured by expression as shown by the colour bar. **(E)** Dotplot visualisation showing scaled gene expression for *in vivo* chondrocytes. Dot size corresponds to the fraction of cells with non-zero expression. Dots are coloured according to gene expression level, according to the adjacent colour key. CC; Chondrocyte **(F)**Uniform Manifold Approximation & Projection (UMAP) plots of day 0, 3, 7 & 14 *in vitro* single nuclei, showing expression of off-target transcripts.Dots are coloured according to gene expression level, according to the adjacent colour key.

**Figure S7.**
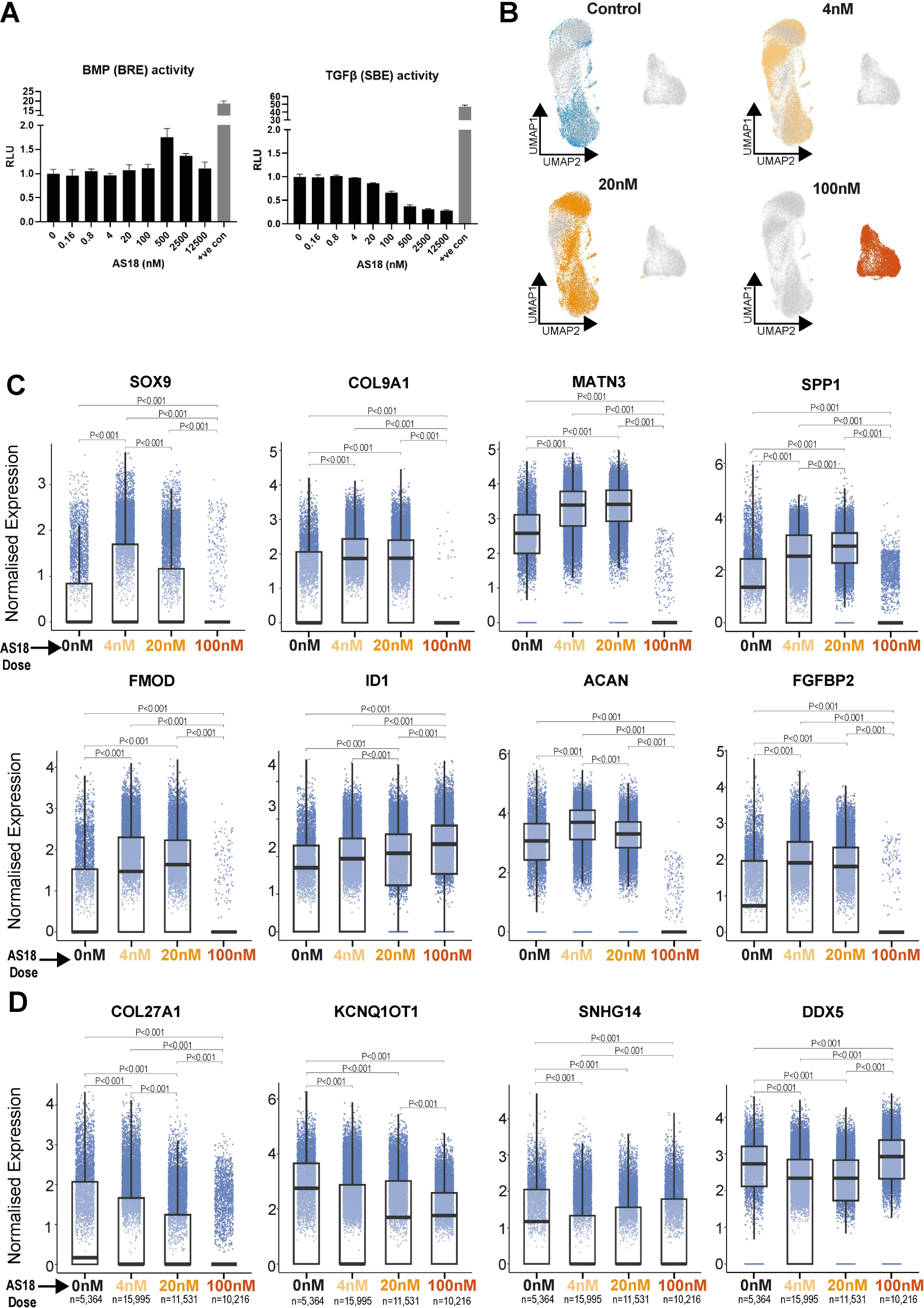
The effect of AS18 inhibition on *in vitro* chondrocytes. **(A)** Bar plots showing reporter cell line activity as varying doses of AS18 **(B)** Uniform Manifold Approximation & Projection (UMAP) plot of day 21 cells treated with 0nM(control), 4nM, 20nM and 100nM of AS18. coloured cells in each panel highlight cells treated with the concentration above that panel. **(C)** Box plots showing the expression of chondrocyte genes at different AS18 concentrations. The box contains the 25th to 75th percentiles of the data, with the central line denoting the median value. The upper whisker extends from the median to the largest value, no further than the 1.5 * inter-quartile range (IQR). The lower whisker extends from the median to the smallest value, at most 1.5 * IQR. Wilcoxon rank-sum test, two-sided, with adjustment for multiple comparisons. **(D)** Box plots showing the expression of osteoblast genes at different AS18 concentrations. The box contains the 25th to 75th percentiles of the data, with the central line denoting the median value. The upper whisker extends from the median to the largest value, no further than the 1.5 * inter-quartile range (IQR). The lower whisker extends from the median to the smallest value, at most 1.5 * IQR. Wilcoxon rank-sum test, two-sided, with adjustment for multiple comparisons.

## Materials and Methods

### Sample acquisition and ethics

The hind limbs of first-trimester human embryos were obtained from elective terminations under REC 96/085 (East of England - Cambridge Central Research Ethics Committee) Three samples (7, 8 & 9 PCW) were maintained in HypoThermosol at 4°C prior to processing to dissection and dissociation for single cell sequencing. A further sample (6 PCW) was collected for in-situ sequencing.

### Tissue Dissection and dissociation- First trimester fetal bone samples

The femora, tibiae and fibulae were dissected from the fetal hind limbs by a specialist bone and soft tissue pathologist [PB] under a microscope using sterile microsurgical instruments. In the 8 & 9 PCW week samples, the femora were further dissected into proximal and distal halves. In the 9 PCW sample only, the tibia was also dissected into proximal and distal halves. Tissue was digested in 5ml of a 5 µg/ml Liberase TH working solution prepared from Liberase TH powder (Sigma 5401135001) and 1X phosphate-buffered saline (PBS) on a shaking platform (750 rpm) at 37°C for 30 minutes. The tissue was gently agitated using a P1000 pipette after 15 minutes. Five ml of 2% fetal bovine serum (FBS) in PBS was then added to stop the dissociation, prior to second stage digestion with 0.25% trypsin solution for a further 30 minutes at 37°C, with pipette agitation every 5 minutes. Cells were then spun down at 750 g at 4°C for 5 min and resuspended in 50-200 µl of 2% FBS in PBS. Fetal cells were loaded for scRNAseq directly following sample processing.

### Human embryonic stem culture

The human embryonic stem cell (hESC) line ‘Man13’ were cultured as colonies under feeder-free conditions in TeSR-E8 culture medium (StemCell Tech 05990), on 6-well tissue culture plates pre-coated with VTN (GIBCO 14700)^1^. Stem cell colonies were passaged every 7 days and before reaching ∼80% confluence using 0.5 mM EDTA (Thermo 15575-038).

### Human embryonic stem differentiation to iMSCs

Four days prior to starting differentiation human embryonic stem cell colonies were passaged using EDTA and plated to produce ∼10 colonies per well of a VTN-coated 6-well plate. To induce differentiation of hESCs to iMSCs, culture medium was switched from TeSR-E8 to MesenPRO RS medium (Gibco 12746012). MesenPRO RS medium was then replaced every 2days. 7days after switching to MesenPRO RS medium, cells were passaged using TrypLE (Gibco 12604-021) and re-plated to a T75 culture flask precoated in a 0.1% gelatine solution (Sigma G1393). After a further 7 days of growth in MesenPRO RS medium, cells were passaged using TrypLE and replated into T75 tissue culture flasks (no additional coating) at a split ratio of 1:8, this is referred to as P1 iMSC. iMSCs were then maintained in T75s using MesenPRO RS™ medium and passaged at 1:8 split ratio once 80% confluent.

### iMSC differentiation to cartilage pellets

Passage 3 to 5 iMSCs were differentiated to chondrocytes in cartilage pellet culture. iMSCs were dissociated using TrypLE, washed with PBS, counted, and then resuspended at a density of 200,000 cells/ml in chondrogenic medium consisting of; DMEM, L-glutamine, 100 nM dexamethasone (Sigma D4902), 50 µg/ml ASC-2-P (Sigma A8960), 40 µg/ml L-proline (Sigma P5607), 1x ITS+l (Sigma I2521), 10 ng/ml TGFβ3 and 100 ng/ml BMP2. 1 ml of cells (200,000) in chondrogenic medium were then pelleted in each 15 ml centrifuge tube. The cap of each 15 ml centrifuge tube was loosened to allow gas exchange and pellets incubated at 37°C and 5% CO2. After 24 h pellets were agitated to ensure detachment from tube and then unless otherwise stated fed every 3 days with fresh chondrogenic medium until pellets were harvested for RNA at day 14, 21, and28 days post pelleting. For FoxO antagonist AS1842856 (AS18) experiments, cartilage pellets were switched to chondrogenic medium containing either 4 nM, 20 nM, 100 nM or 500 nM AS18 at day 14, which was then replaced at day 17 and then pellets harvested for RNA at day 21. RNA was extracted using molecular grinding resin with pestles and the QIAGEN RNeasy Mini Kit.

### *In vitro* tissue RNA isolation and sample preparation for bulk RNA sequencing

RNA samples were isolated using miRvana microRNA Isolation Kit (Thermo). Total RNA was collected at day 0 (stage 0 samples), day 3 (stage 2 samples), day 7, day 14 and day 21 of the protocol. A total of 1 µg of RNA was used for library preparation using the Illumina TruSeq Stranded mRNA Sample preparation kit for transcriptome libraries and Illumina TruSeq Small RNA Library Preparation kit for small RNA libraries. All samples were run on an Illumina HiSeq 2000 with paired-end reads, generating on average 50 million high-quality sequencing reads per sample.

### Bulk RNA sequencing of in vitro tissue

Assessment of raw read quality was performed using FastQC v0.11.6. Paired-end reads were aligned using STAR 2.7.7a ^2^ to the primary assembly of the human genome (version GRCh38.p5), with GENCODE v26 annotation ^3^. ENCODE STAR options were enabled, thus generating alignments to both genome and transcriptome. Genome BAM files were used to generate TDF files using igvtools v2.3.93, and visualised using IGV v2.4.11 ^4^. Transcriptomic BAM files were used for quantification with RSEM v1.3.3 ^5^. This generated expression tables in raw counts, TPM, and FPKM, summarised on a transcript and gene level that were used for further analysis.

Raw data was then pre-processed using the edgeR package ^6^. First, raw count data were converted to log-CPM values, and reads with low expression were filtered out to keep only those with at least ten counts in three samples or more. Normalisation was then applied using the trimmed mean of M-values (TMM) method. Visualisation of gene expression was performed using the ComplexHeatmap package for R^7^.

### *In vitro* tissue processing for single-nuclei RNA sequencing

Frozen tissue samples were cryosectioned into 10 µm sections before being transferred to homogenisation buffer in a dounce homogeniser for initial dissociation through 20 strokes with a loose pestle, then a further 20 strokes with a tight pestle, followed by visual confirmation that the tissue had dissociated. Filtration through a 40 µm cell strainer on ice was performed to ensure no larger fragments remained, and visual inspection of the remaining nuclei under microscope was performed to ensure no clumping had occurred. If clumping was present an additional Percoll clean-up step was carried out. Nuclei were suspended in wash buffer and counted manually using a C-chip with Trypan blue staining.

### Single-cell & single-nuclei encapsulation and RNA sequencing

Single-cell/ single-nuclei suspensions were loaded onto individual channels of a Chromium 10x Genomics single cell 3’ v2 library chip as per the manufacturer’s protocol (10x Genomics; PN-120233). cDNA sequencing libraries were prepared as per the manufacturer’s protocol and sequenced using an Illumina HiSeq 4000 with 2×50bp paired-end reads.

### scRNA-seq alignment, processing and annotation

Newly generated and publicly available raw 10X sequencing data were aligned and quantified using starsolo 2.7.9a^8^. Cells were called using EmptyDrops^9^. The human reference hg38 genome refdata-gex-GRCh38-2020-A was utilised, available at: https://cf.10xgenomics.com/supp/cell-exp/refdata-gex-GRCh38-2020-A.tar.gz.

Cell-free RNAs were removed using SoupX1.4.5 with default parameters^10^. Cells that expressed fewer than 500 genes or more than 7,500 were excluded, as were cells where mitochondrial transcripts accounted for >10% of the overall transcriptome. After normalising each cell against total UMI counts and log transformation using Scanpy1.8.2, highly variable genes were calculated using Scanpy’s default ^11^. The feature gene count matrix was scaled across cells for each gene and fed into Principal Component Analysis (PCA). For fetal limb data, BBKNN was then used to integrate newly generated and publicly available single cell datasets together (batch_key=’batch’, neighbors_within_batch = 3, trim = 200,approx=False,metric=’euclidean’,n_pcs=100)^12^. The resulting AnnData object was converted for use with the Seurat version 4.0.1 package for R, using the “convertFormat” function of the sceasy R package ^13^. Differential gene expression testing between clusters was performed using the Wilcoxon rank sum test corrected for multiple comparisons, within Seurat’s ‘FindMarkers’ function, using default parameters, and clusters annotated based on classical marker genes. Gene expression was visualised using the “DotPlot” and “FeaturePlot” Seurat functions as well as the ggplot2 package for R^14^. For constructing a heatmap of marker genes, the “DoHeatmap” function was used, after subsetting each celltype to 250 cells, or the total number captured if less than 250.

### Doublet detection

Expression matrices were processed with the Scrublet V0.2.2 pipeline using default parameters^15^. Clustering was then re-run at resolution 10 to provide an over-clustered manifold, and any clusters in which more than 20% of cells were called as doublets were removed (a single cluster of 155 cells, of which 70 were called as doublets). Subsequently, any remaining cells called as doublets were removed from analysis. A total of 828 cells were removed from fetal limb data, and 76 from *in vitro* data.

### Pseudotime Computation

The pseudotime of *in vivo* cells was estimated by running the Gaussian Process Latent Variable Model (GPLVM) pseudotime estimator implemented in Pyro^16^. Root state and prior cell type ordering was based on annotated cell type. Pseudotime was visualised using the sc.pl.umap command within scanpy, colouring by GPLVM pseudotime.

### Differential abundance testing

In order to examine the relative abundance of chondrocyte celltypes in different samples along the proximo-distal axis of the limb, we applied the MiloR package^17^. The K-nearest neighbour graph was first constructed using the “buildGraph” function with default parameters, and neighbourhoods defined with the “makeNhoods” function, with 10% of total cells randomly selected for construction. Neighbourhood testing was performed to test for differential cell type abundance in the femur versus tibia & fibula, accounting for PCW of the sample as a covariate. Results were visualised using the “plotDAbeeswarm” function in MiloR. There were insufficient hypertrophic chondrocytes to calculate adequately powered abundances.

### Transcription factor network inference

Regulon activity within bulk and single-cell sequencing data were computed using the DoRothEA package for R ^18,19^. The activity of regulons within the bulk or single cell datasets was calculated using the wrapper for the Viper package for R contained within DoRothEA, using the “run_viper” function^20^. Activity scores per cluster or bulk sample were then visualised using the ComplexHeatmap package for R^7^.

### Fetal single-cell to in vitro bulk data comparison

The cell-type annotated fetal single cell data was used as a reference map to determine the composition of the *in vitro* tissue. The abundance of fetal cellular signals in the *In vitro* bulk transcriptomes was quantified using the CellSignalAnalysis python package and visualised using the ComplexHeatmap package for R ^7,21^. Genes in the non-shared space were dropped prior to analysis.

### *In vitro* and *in vivo* trajectory comparison

To understand the agreement and differences between *in vitro* and *in vivo* chondrocyte differentiation, we performed a computational alignment between them using dynamic programming (DP). Genes2Genes (G2G) ^22^ is a Bayesian information-theoretic DP framework that quantifies the similarity between two single-cell transcriptomic pseudotime trajectories by generating their optimal alignment at both gene-level and cell-level. A pseudotime trajectory alignment describes a non-linear mapping between the *in vitro* and *in vivo* pseudo timepoints in sequential order. This mapping may include three possible mappings between a pair of *in vitro* - *in vivo* time points, i.e., one-to-one match (denoted by M), one-to-many match (where one point in a reference pseudotime matches to multiple points in query pseudotime, denoted by V), and many-to-one match (where multiple points in query pseudotime match to a single point in reference pseudotime, denoted by W). Similarly, the mapping can detect any mismatched time points, i.e. insertion (where a time point in the query pseudotime has no corresponding timepoint in the reference pseudotime, denoted by I) and deletion (the inverse of insertion, denoted by D. Accordingly, for each gene of interest, G2G outputs a 5-state alignment string, defining the optimal sequence of matched and mismatched time points between them (which is analogous to an output from DNA/protein sequence alignment).

To generate such alignment, G2G optimises the cost of matching (i.e. null hypothesis) or mismatching (i.e. alternative hypothesis) the gene expression distributions of each pair of *in vitro - in vivo* time points, based on an entropy difference (a Shannon information distance measured in the unit of information *nits*) between the two hypotheses under the Minimum Message Length criterion^23^. Each gene alignment carries an alignment similarity measure (i.e. the percentage of matches across the alignment string). G2G also generates an aggregated cell-level alignment and similarity percentage statistic by averaging across all gene alignments.

#### Preparing in vivo, in vitro trajectories for comparison

Prior to trajectory alignment, we obtained the analogous chondrocyte trajectories for comparison by removing the non-chondrocyte lineage cells from the *in vivo* data, leaving mesenchymal condensate and chondrocyte cell types. In order to have the two trajectories end at a comparable point of chondrocyte maturation and prevent over-representation of the hypertrophic chondrocyte state in the *in vitro* trajectory, day 21 data was not included. This resulted in 22,181 *in vitro* cells. We also downsampled the *in vivo* dataset by subsampling (i.e. uniform random sampling) from each different cell type aiming for ∼20,000 cells, to have the number of *in vivo* cells approximately similar to that of the *in vitro* data. If a cell type had less than 500 cells, we retained all cells. This reduced *in vivo* representation resulted in 20,041 *in vivo* cells.

Next we estimated the pseudotime of *in vivo* and *in vitro* cells by running the Gaussian Process Latent Variable Model (GPLVM) pseudotime estimator implemented in Pyro^16^. GPLVM has shown to be successful in incorporating useful time priors for single-cell trajectory inference. For *in vivo* pseduotime estimation, we ordered the cells based on their annotated cell type and their known order in the lineage, to give them as priors for the GPLVM. For *in vitro* pseduotime estimation, we used their capture day time as priors for GPLVM. The resultant pseudotime estimates were min-max normalised before alignment. The optimal binning of the pseudotime axis was inferred using the *optBinning* python package ^24^ based on the pseudotime distributions, resulting in 12 optimal time bins.

#### Computational Alignment

Given the normalised gene expression data (log1p normalised after normalising for total transcript count to add up to 10,000), and their GPLVM pseudotime estimates, we ran alignment for 275 genes (after choosing only the highly variable genes and filtering out ones with less than 10 cells expressed). G2G generated fully descriptive 5-state alignment strings for each gene along with their alignment similarity percentage statistics, as well as an average alignment across all those genes. We also ran the alignment for 288 transcription factors (TFs) (chosen from the list in ^25^ and by performing the same filtering as before). All results were visualised using the plotting functions of G2G that are based on standard *Matplotlib* and *Seaborn* Python libraries.

### In vitro AS18 inhibition differential expression testing

Day 21 cells were grouped based on AS18 dose at day 14 (0 nM/control, 4 nM, 20 nM & 100 nM) and differential gene expression testing was performed using the Wilcoxon rank sum test corrected for multiple comparisons, within Seurat’s ‘FindMarkers’ function, using default parameters. The two groups being compared were specified using the “ident.1” and “ident.2” parameters of the test. Expression was visualised with boxplots, using the ggplot2 package for R^14^. For analysis of apoptosis genes in the 100nM treated *in vitro* cells, the apoptosis gene set was extracted from the Kyoto Encyclopedia of Genes and Genomes (KEGG) and cross referenced against the significantly differentially expressed genes in 100nM cells ^26^.

### CARTANA in situ sequencing

The lower limb of a 7.5 GW human embryo was embedded in optimal cutting temperature medium (OCT) and frozen at −80°C on an isopentane-dry ice slurry. Cryosections were cut at a thickness of 10 μm using a Leica CM1950 cryostat and placed onto SuperFrost Plus slides (VWR).

In situ sequencing was performed using the 10X Genomics CARTANA HS Library Preparation Kit (1110-02, following protocol D025) and In Situ Sequencing Kit (3110-02, following protocol D100), which comprise a commercialised version of HybRISS^27^.

Briefly: a limb section was fixed in 3.7% formaldehyde (Merck 252549) in PBS for 30 minutes, washed twice in PBS for 1 minute each, permeabilised in 0.1 M HCl (Fisher 10325710) for 5 minutes, and washed twice again in PBS, all at room temperature. Following dehydration in 70% and 100% ethanol for 2 minutes each, a 9 mm diameter (50 μl volume) SecureSeal hybridisation chamber (Grace Bio-Labs GBL621505-20EA) was adhered to the slide and used to hold subsequent reaction mixtures. Following rehydration in buffer WB3, probe hybridisation in buffer RM1 was conducted for 16 hours at 37°C. The 90-plex probe panel included 5 padlock probes per gene, the sequences of which are proprietary (10X Genomics CARTANA). The section was washed with PBS-T (PBS with 0.05% Tween-20) twice, then with buffer WB4 for 30 minutes at 37°C, and thrice again with PBS-T. Probe ligation in RM2 was conducted for 2 hours at 37°C and the section washed thrice with PBS-T, then rolling circle amplification in RM3 was conducted for 18 hours at 30°C. Following PBS-T washes, all rolling circle products (RCPs) were hybridised with LM (Cy5 labelling mix with DAPI) for 30 minutes at room temperature, the section was washed with PBS-T and dehydrated with 70% and 100% ethanol. The hybridisation chamber was removed and the slide mounted with SlowFade Gold Antifade Mountant (Thermo S36937). Imaging of Cy5-labelled RCPs at this stage acted as a QC step to confirm RCP (‘anchor’) generation and served to identify spots during decoding. Imaging was conducted using a Perkin Elmer Opera Phenix Plus High-Content Screening System in confocal mode with 1 μm z-step size, using a 63X (NA 1.15, 0.097 μm/pixel) water-immersion objective. Channels: DAPI (excitation 375 nm, emission 435-480 nm), Atto 425 (ex. 425 nm, em. 463-501 nm), Alexa Fluor 488 (ex. 488 nm, em. 500-550 nm), Cy3 (ex. 561 nm, em. 570-630 nm), Cy5 (ex. 640 nm, em. 650-760 nm).

Following imaging, the slide was de-coverslipped vertically in PBS (gently, with minimal agitation such that the coverslip ‘fell’ off to prevent damage to the tissue). The section was dehydrated with 70% and 100% ethanol, and a new hybridisation chamber secured to the slide. The previous cycle was stripped using 100% formamide (Thermo AM9342), which was applied fresh each minute for 5 minutes, then washed with PBS-T. Barcode labelling was conducted using two rounds of hybridisation, first an adapter probe pool (AP mixes AP1-AP6, in subsequent cycles), then a sequencing pool (SP mix with DAPI, customised with Atto 425 in place of Alexa Fluor 750), each for 1 hour at 37°C with PBS-T washes in between and after. The section was dehydrated, the chamber removed, and the slide mounted and imaged as previously. This was repeated another five times to generate the full dataset of 7 cycles (anchor and 6 barcode bits).

#### Lower limb in-situ sequencing (ISS) image data processing

##### 1. Image stitching

The raw data obtained from the microscope underwent initial processing by utilising proprietary software provided by Perkin Elmer. This entailed maximum Z intensity projection and stitching procedures using the 7% overlap between adjacent tiles, resulting in the generation of an ome.tiff file per imaging cycle that encompasses all the channels (DAPI, Atto425, Alexa Fluor 488, Cy3 and Cy5).

##### 2. Cyclewise image registration

The cycle-wise image registration is done in two steps using the Microaligner package ^28,29^. The first one is the Affine feature-based registration. This method is an image registration algorithm that begins by detecting image features in the DAPI image of the first imaging cycle using the FAST feature finder algorithm, which identifies areas with significant intensity changes. Next, the DAISY feature descriptor algorithm extracts histograms of oriented gradients for each identified feature. The extracted features are then matched using the FLANN-based KNN matcher algorithm, which determines the correspondence between the features of the reference and moving images. The matches are filtered based on the default distance threshold between neighbouring features, and the resulting matched feature coordinates are aligned using the RANSAC algorithm to compute the affine transformation. This initial phase aims to align images with significant linear drifts across cycles. The process involves the partitioning of images into tiles of 6000 by 6000 pixels to optimise the alignment and reduce memory usage. For each tile, a transformation matrix is derived after applying the DoG function with predefined kernel sizes, which is eventually unified by employing the matmul function in Python.

The second step non-linear optical flow-based registration, in contrast, relies on the Farneback method (calcOpticalFlowFarneback) in OpenCV with fine tuned parameters - pyr_scale=0.5, levels=3, winsize=51, iterations=3, poly_n=1, poly_sigma=1.7, flags=OPTFLOW_FARNEBACK_GAUSSIAN - which identifies pixels with the highest similarity within a given window. For each pixel, the method computes a 2D vector that characterises the movement of the pixel from one image to the other. This approach is particularly suitable for aligning small local shifts across images and as the first step is applied tile by tile.

##### 3. ISS barcode decoding with PoSTcode

To decode individual RNA transcripts from cyclic ISS images, we used the PoSTcode barcode decoding algorithm ^28^ and customised image preprocessing.To improve the accuracy of downstream spot-calling and quality of intensity-based decoding, we first applied the white hat filter with the kernel size of 5 pixels to filter out noise from all coding channels. Subsequently, the transcript detection was performed exclusively on the anchor channel using the ‘locate’ method in TrackPy ^30^ with percentile equal 90, spot size equals 5 and separation equals 4. One intensity profile for each transcript was extracted from the registered image generated in the last step. This intensity profile is of shape 4×6 representing the intensity extracted from 4 channels per cycle and from all the 6 cycles. Yet, to improve decoding outcome, we expanded the searching range of the maximum intensity to +/− 2 pixels across coding channels. The decoding step in PoSTcode takes this 4×6 matrix and the codebook from CARTANA as input and returns prediction of gene type for each transcript with a confidence value. Only transcripts with value higher than 0.97 were kept and saved as a .csv file for downstream processing.

##### 4. Single cell segmentation

To segment single cells from the registered image stack, we applied the cell segmentation in CellPose ^31^ on DAPI channel with the cell size of 70 pixels in diameter. Due to the large memory requirements, we adopted a strategy of dividing the whole images into smaller tiles and performed the segmentation on each of the tiles individually. Following this, we stitched the tiles back together to reconstruct the complete image without compromising much segmentation accuracy. There were in total 117,788 cells detected.

##### 5. Anndata object generation

The decoded 1,164,802 spots were assigned to the 117,788 cells using the STRtree in ^32^. Out of the 117,788 cells, only 66,675 cells were kept after filtering out cells with less than 4 transcripts. The output is saved as an anndata object in h5ad format.

### Imputation of single cell locations in ISS data

Approximate expression signatures for genes missing in the ISS data were constructed based on matched scRNA-Seq data. The 90 genes present in the ISS pool were isolated, and the scRNA-Seq and ISS data were separately normalised to median total cell counts per modality and log-transformed. The 15 nearest neighbours in scRNA-Seq space were identified for each ISS cell with annoy, and the genes absent from ISS were imputed as the means of the raw counts of the scRNA-seq neighbours.

### Multiplexed smFISH (RNAscope)

Cryosections were processed using a Leica BOND RX to automate staining with the RNAscope Multiplex Fluorescent Reagent Kit v2 Assay (Advanced Cell Diagnostics, Bio-Techne), according to the manufacturers’ instructions. Prior to staining, fresh frozen sections were post-fixed in 4% paraformaldehyde in PBS for 15 minutes at 4°C, then dehydrated through a series of 50%, 70%, 100%, and 100% ethanol, for 5 minutes each. Following manual pre-treatment, automated processing included digestion with Protease IV for 30 minutes prior to probe hybridisation. Tyramide signal amplification with Opal 520, Opal 570, and Opal 650 and TSA-biotin (TSA Plus Biotin Kit) (Akoya Biosciences) and streptavidin-conjugated Atto 425 (Sigma) was used to develop RNAscope probe channels. Stained sections were imaged using a Perkin Elmer Opera Phenix Plus High-Content Screening System in confocal mode with 1 μm z-step size, using a 63X (NA 1.15, 0.097 μm/pixel) water-immersion objective. Channels: DAPI (excitation 375 nm, emission 435-480 nm), Atto 425 (ex. 425 nm, em. 463-501 nm), Opal 520 (ex. 488 nm, em. 500-550 nm), Opal 570 (ex. 561 nm, em. 570-630 nm), Opal 650 (ex. 640 nm, em. 650-760 nm).

## Acknowledgements

We thank Professor Matt Hurles, Dr Emma Davenport, Professor Matthew Allen and Dr Mekayla Storer for their valuable feedback on the study.

## Funding

This study was supported by the Wellcome Trust through institutional / programme grants and personal Fellowships (222902/Z/21/Z, 206194, 220540/Z/20/A and 211276/Z/18/Z).

## Author contributions

Study conception: J.E.L., S.W., K.R., S.A.T, S.K, A.F., S.B. Data analysis and interpretation: J.E.L, S.W, K.R, D.S, T.L, A.V.P, P.H, K.P, S.W, L.J, M.J.Y, S.B, O.B, M.H. Experiments: K.R., E.T., P.B, J.E.L., E.P., D.Z., L.M. Tissue curation: X.H, R.B. Manuscript writing: J.E.L, S.W, K.R, S.K, S.A.T with contributions from all authors. S.A.T. and S.K. co-directed the study.

## Competing Interests

In the past three years, S.A.T. has consulted for or been a member of scientific advisory boards at Qiagen, Sanofi, GlaxoSmithKline and ForeSite Labs. She is a consultant and equity holder for TransitionBio and EnsoCell. The remaining authors declare no competing interests.

## Data Availability

All of our newly generated raw single-cell RNA sequencing data from embryonic hindlimb long bones and *in vitro* chondrogenesis protocols are available via the European Genome-Phenome archive via the accession number EGAD00001011306.

Previously published raw data utilised in this study are available from ArrayExpress (first trimester whole-limb samples E-MTAB-8813; second trimester bone marrow samples E-MTAB-9389). Processed data can be visualised at our portal: https://chondrocytes.cellgeni.sanger.ac.uk/

## Code availability

No previously unreported custom computer code or algorithms were used to generate results that are reported in this paper

## Extended Data

Supplementary figures S1 to 7

Supplementary tables 1 to 6

## Notes

https://chondrocytes.cellgeni.sanger.ac.uk/

## References

1. Aldridge, S., and Teichmann, S.A. (2020). Single cell transcriptomics comes of age. Nat. Commun. 11. 10.1038/s41467-020-18158-5.

2. Behjati, S., Lindsay, S., Teichmann, S.A., and Haniffa, M. (2018). Mapping human development at single-cell resolution. Development 145. 10.1242/dev.152561.

3. Rood, J.E., Maartens, A., Hupalowska, A., Teichmann, S.A., and Regev, A. (2022). Impact of the Human Cell Atlas on medicine. Nat. Med. 28, 2486–2496.

4. Garcia-Alonso, L., Handfield, L.-F., Roberts, K., Nikolakopoulou, K., Fernando, R.C., Gardner, L., Woodhams, B., Arutyunyan, A., Polanski, K., Hoo, R., et al. (2021). Mapping the temporal and spatial dynamics of the human endometrium in vivo and in vitro. Nat. Genet. 53, 1698–1711.

5. Fujii, M., Matano, M., Toshimitsu, K., Takano, A., Mikami, Y., Nishikori, S., Sugimoto, S., and Sato, T. (2018). Human Intestinal Organoids Maintain Self-Renewal Capacity and Cellular Diversity in Niche-Inspired Culture Condition. Cell Stem Cell 23. 10.1016/j.stem.2018.11.016.

6. GBD 2019 Diseases and Injuries Collaborators (2020). Global burden of 369 diseases and injuries in 204 countries and territories, 1990-2019: a systematic analysis for the Global Burden of Disease Study 2019. Lancet 396, 1204–1222.

7. Ackerman, I.N., Bohensky, M.A., Zomer, E., Tacey, M., Gorelik, A., Brand, C.A., and de Steiger, R. (2019). The projected burden of primary total knee and hip replacement for osteoarthritis in Australia to the year 2030. BMC Musculoskelet. Disord. 20, 90.

8. Roseti, L., Desando, G., Cavallo, C., Petretta, M., and Grigolo, B. (2019). Articular Cartilage Regeneration in Osteoarthritis. Cells 8. 10.3390/cells8111305.

9. Augustyniak, E., Trzeciak, T., Richter, M., Kaczmarczyk, J., and Suchorska, W. (2015). The role of growth factors in stem cell-directed chondrogenesis: a real hope for damaged cartilage regeneration. Int. Orthop. 39. 10.1007/s00264-014-2619-0.

10. Andreas, K., Lübke, C., Häupl, T., Dehne, T., Morawietz, L., Ringe, J., Kaps, C., and Sittinger, M. (2008). Key regulatory molecules of cartilage destruction in rheumatoid arthritis: an in vitro study. Arthritis Res. Ther. 10, 1–16.

11. Kim, H.J., and Park, J.S. (2017). Usage of Human Mesenchymal Stem Cells in Cell-based Therapy: Advantages and Disadvantages. Development & reproduction 21. 10.12717/DR.2017.21.1.001.

12. Karlsen, T.A., Sundaram, A.Y.M., and Brinchmann, J.E. (2019). Single-Cell RNA Sequencing of In Vitro Expanded Chondrocytes: MSC-Like Cells With No Evidence of Distinct Subsets. Cartilage. 10.1177/1947603519847746.

13. Manolagas, S.C., and Kronenberg, H.M. (2014). Reproducibility of results in preclinical studies: a perspective from the bone field. J. Bone Miner. Res. 29. 10.1002/jbmr.2293.

14. Suchorska, W.M., Augustyniak, E., Richter, M., and Trzeciak, T. (2017). Comparison of Four Protocols to Generate Chondrocyte-Like Cells from Human Induced Pluripotent Stem Cells (hiPSCs). Stem cell reviews and reports 13. 10.1007/s12015-016-9708-y.

15. Endochondral ossification: How cartilage is converted into bone in the developing skeleton (2008). Int. J. Biochem. Cell Biol. 40, 46–62.

16. Larry Jameson, J., and De Groot, L.J. (2015). Endocrinology: Adult and Pediatric E-Book (Elsevier Health Sciences).

17. Shwartz, Y., Viukov, S., Krief, S., and Zelzer, E. (2016). Joint Development Involves a Continuous Influx of Gdf5-Positive Cells. Cell Rep. 15, 2577.

18. Bian, Q., Cheng, Y.-H., Wilson, J.P., Su, E.Y., Kim, D.W., Wang, H., Yoo, S., Blackshaw, S., and Cahan, P. (2020). A single cell transcriptional atlas of early synovial joint development. Development 147. 10.1242/dev.185777.

19. Zhang, B., He, P., Lawrence, J.E., Wang, S., Tuck, E., Williams, B., Roberts, K., Kleshchevnikov, V., Mamanova, L., Bolt, L., et al. (2022). A human embryonic limb cell atlas resolved in space and time. bioRxiv, 2022.04.27.489800. 10.1101/2022.04.27.489800.

20. Jardine, L., Webb, S., Goh, I., Londoño, M.Q., Reynolds, G., Mather, M., Olabi, B., Stephenson, E., Botting, R.A., Horsfall, D., et al. (2021). Blood and immune development in human fetal bone marrow and Down syndrome. Nature 598, 327.

21. Li, T., Horsfall, D., Basurto-Lozada, D., Roberts, K., Prete, M., Lawrence, J.E.G., He, P., Tuck, E., Moore, J., Ghazanfar, S., et al. (2023). WebAtlas pipeline for integrated single cell and spatial transcriptomic data. bioRxiv, 2023.05.19.541329. 10.1101/2023.05.19.541329.

22. Ma, X., Su, P., Yin, C., Lin, X., Wang, X., Gao, Y., Patil, S., War, A.R., Qadir, A., Tian, Y., et al. (2020). The Roles of FoxO Transcription Factors in Regulation of Bone Cells Function. Int. J. Mol. Sci. 21, 692.

23. Yamada, D., Nakamura, M., Takao, T., Takihira, S., Yoshida, A., Kawai, S., Miura, A., Ming, L., Yoshitomi, H., Gozu, M., et al. (2021). Induction and expansion of human PRRX1+ limb-bud-like mesenchymal cells from pluripotent stem cells. Nature biomedical engineering 5. 10.1038/s41551-021-00778-x.

24. Guzzo, R.M., Andreeva, V., Spicer, D.B., and Drissi, M.H. (2011). Persistent expression of Twist1 in chondrocytes causes growth plate abnormalities and dwarfism in mice. Int. J. Dev. Biol. 55. 10.1387/ijdb.103274rg.

25. Reinhold, M.I., Kapadia, R.M., Liao, Z., and Naski, M.C. (2006). The Wnt-inducible transcription factor Twist1 inhibits chondrogenesis. J. Biol. Chem. 281. 10.1074/jbc.M504875200.

26. Zhao, H., Zhou, W., Yao, Z., Wan, Y., Cao, J., Zhang, L., Zhao, J., Li, H., Zhou, R., Li, B., et al. (2015). Foxp1/2/4 regulate endochondral ossification as a suppresser complex. Dev. Biol. 398, 242.

27. Hallett, S.A., Matsushita, Y., Ono, W., Sakagami, N., Mizuhashi, K., Tokavanich, N., Nagata, M., Zhou, A., Hirai, T., Kronenberg, H.M., et al. (2021). Chondrocytes in the resting zone of the growth plate are maintained in a Wnt-inhibitory environment. 10.7554/eLife.64513.

28. Surmann-Schmitt, C., Dietz, U., Kireva, T., Adam, N., Park, J., Tagariello, A., Onnerfjord, P., Heinegård, D., Schlötzer-Schrehardt, U., Deutzmann, R., et al. (2008). Ucma, a novel secreted cartilage-specific protein with implications in osteogenesis. J. Biol. Chem. 283. 10.1074/jbc.M702792200.

29. St-Jacques, B., Hammerschmidt, M., and McMahon, A.P. (1999). Indian hedgehog signaling regulates proliferation and differentiation of chondrocytes and is essential for bone formation. Genes Dev. 13. 10.1101/gad.13.16.2072.

30. Kielty, C.M., Kwan, A.P., Holmes, D.F., Schor, S.L., and Grant, M.E. (1985). Type X collagen, a product of hypertrophic chondrocytes. Biochem. J. 227. 10.1042/bj2270545.

31. Cancela, M.L., Conceição, N., and Laizé, V. (2012). Gla-rich protein, a new player in tissue calcification? Adv. Nutr. 3, 174–181.

32. Kozhemyakina, E., Zhang, M., Ionescu, A., Ayturk, U.M., Ono, N., Kobayashi, A., Kronenberg, H., Warman, M.L., and Lassar, A.B. (2015). Identification of a Prg4-expressing articular cartilage progenitor cell population in mice. Arthritis & rheumatology (Hoboken, N.J.) 67. 10.1002/art.39030.

33. Mobasheri, A., Trujillo, E., Bell, S., Carter, S.D., Clegg, P.D., Martín-Vasallo, P., and Marples, D. (2004). Aquaporin water channels AQP1 and AQP3, are expressed in equine articular chondrocytes. Vet. J. 168. 10.1016/j.tvjl.2003.08.001.

34. Merino, R., Macias, D., Gañan, Y., Economides, A.N., Wang, X., Wu, Q., Stahl, N., Sampath, K.T., Varona, P., and Hurle, J.M. (1999). Expression and function of Gdf-5 during digit skeletogenesis in the embryonic chick leg bud. Dev. Biol. 206. 10.1006/dbio.1998.9129.

35. Bandyopadhyay, A., Kubilus, J.K., Crochiere, M.L., Linsenmayer, T.F., and Tabin, C.J. (2008). Identification of unique molecular subdomains in the perichondrium and periosteum and their role in regulating gene expression in the underlying chondrocytes. Dev. Biol. 321, 162.

36. Bialek, P., Kern, B., Yang, X., Schrock, M., Sosic, D., Hong, N., Wu, H., Yu, K., Ornitz, D.M., Olson, E.N., et al. (2004). A twist code determines the onset of osteoblast differentiation. Dev. Cell 6, 423–435.

37. Nagano, K. (2019). R-spondin signaling as a pivotal regulator of tissue development and homeostasis. Jpn. Dent. Sci. Rev. 55. 10.1016/j.jdsr.2019.03.001.

38. Bei, K., Du, Z., Xiong, Y., Liao, J., Su, B., and Wu, L. (2012). BMP7 can promote osteogenic differentiation of human periosteal cells in vitro. Mol. Biol. Rep. 39. 10.1007/s11033-012-1748-z.

39. Lin, W., Zhu, X., Gao, L., Mao, M., Gao, D., and Huang, Z. (2021). Osteomodulin positively regulates osteogenesis through interaction with BMP2. Cell Death Dis. 12, 1–13.

40. Engsig, M.T., Chen, Q.-J., Vu, T.H., Pedersen, A.-C., Therkidsen, B., Lund, L.R., Henriksen, K., Lenhard, T., Foged, N.T., Werb, Z., et al. (2000). Matrix Metalloproteinase 9 and Vascular Endothelial Growth Factor Are Essential for Osteoclast Recruitment into Developing Long Bones. J. Cell Biol. 151, 879.

41. Karsenty, G., Kronenberg, H.M., and Settembre, C. (2009). Genetic control of bone formation. Annu. Rev. Cell Dev. Biol. 25. 10.1146/annurev.cellbio.042308.113308.

42. Matsumoto, T., Yamada, A., Aizawa, R., Suzuki, D., Tsukasaki, M., Suzuki, W., Nakayama, M., Maki, K., Yamamoto, M., Baba, K., et al. (2013). BMP-2 Induced Expression of Alx3 That Is a Positive Regulator of Osteoblast Differentiation. PLoS One 8. 10.1371/journal.pone.0068774.

43. Salie, R., Kneissel, M., Vukevic, M., Zamurovic, N., Kramer, I., Evans, G., Gerwin, N., Mueller, M., Kinzel, B., and Susa, M. (2010). Ubiquitous overexpression of Hey1 transcription factor leads to osteopenia and chondrocyte hypertrophy in bone. Bone 46. 10.1016/j.bone.2009.10.022.

44. Rutkovskiy, A., Stensløkken, K.-O., and Vaage, I.J. (2016). Osteoblast Differentiation at a Glance. Med. Sci. Monit. Basic Res. 22, 95.

45. Youlten, S.E., Kemp, J.P., Logan, J.G., Ghirardello, E.J., Sergio, C.M., Dack, M.R.G., Guilfoyle, S.E., Leitch, V.D., Butterfield, N.C., Komla-Ebri, D., et al. (2021). Osteocyte transcriptome mapping identifies a molecular landscape controlling skeletal homeostasis and susceptibility to skeletal disease. Nat. Commun. 12, 1–21.

46. Minkin, C. (1982). Bone acid phosphatase: tartrate-resistant acid phosphatase as a marker of osteoclast function. Calcif. Tissue Int. 34. 10.1007/BF02411252.

47. Sundaram, K., Nishimura, R., Senn, J., Youssef, R.F., London, S.D., and Reddy, S.V. (2007). RANK ligand signaling modulates the matrix metalloproteinase-9 gene expression during osteoclast differentiation. Exp. Cell Res. 313. 10.1016/j.yexcr.2006.10.001.

48. Matsushita, Y., Ono, W., and Ono, N. (2022). Toward Marrow Adipocytes: Adipogenic Trajectory of the Bone Marrow Stromal Cell Lineage. Front. Endocrinol. 13. 10.3389/fendo.2022.882297.

49. Xie, M., Kamenev, D., Kaucka, M., Kastriti, M.E., Zhou, B., Artemov, A.V., Storer, M., Fried, K., Adameyko, I., Dyachuk, V., et al. (2019). Schwann cell precursors contribute to skeletal formation during embryonic development in mice and zebrafish. Proc. Natl. Acad. Sci. U. S. A. 116. 10.1073/pnas.1900038116.

50. Fountain, D.M., and Sauka-Spengler, T. (2023). The SWI/SNF Complex in Neural Crest Cell Development and Disease. Annu. Rev. Genomics Hum. Genet. 24. 10.1146/annurev-genom-011723-082913.

51. Garcia-Alonso, L., Holland, C.H., Ibrahim, M.M., Turei, D., and Saez-Rodriguez, J. (2019). Benchmark and integration of resources for the estimation of human transcription factor activities. Genome Res. 29, 1363–1375.

52. Holland, C.H., Tanevski, J., Perales-Patón, J., Gleixner, J., Kumar, M.P., Mereu, E., Joughin, B.A., Stegle, O., Lauffenburger, D.A., Heyn, H., et al. (2020). Robustness and applicability of transcription factor and pathway analysis tools on single-cell RNA-seq data. Genome Biol. 21, 36.

53. Liu, C.-F., and Lefebvre, V. (2015). The transcription factors SOX9 and SOX5/SOX6 cooperate genome-wide through super-enhancers to drive chondrogenesis. Nucleic Acids Res. 43, 8183–8203.

54. Fujikawa, J., Takeuchi, Y., Kanazawa, S., Nomir, A.G., Kito, A., Elkhashab, E., Ghaleb, A.M., Yang, V.W., Akiyama, S., Morisaki, I., et al. (2017). Kruppel-like factor 4 regulates matrix metalloproteinase and aggrecanase gene expression in chondrocytes. Cell Tissue Res. 370, 441–449.

55. Macdonald, C.D., Falconer, A.M.D., Chan, C.M., Wilkinson, D.J., Skelton, A., Reynard, L., Litherland, G.J., Nicholas Europe-Finner, G., and Rowan, A.D. (2018). Cytokine-induced cysteine-serine-rich nuclear protein-1 (CSRNP1) selectively contributes to MMP1 expression in human chondrocytes. PLoS One 13. 10.1371/journal.pone.0207240.

56. Liu, N.Q., Lin, Y., Li, L., Lu, J., Geng, D., Zhang, J., Jashashvili, T., Buser, Z., Magallanes, J., Tassey, J., et al. (2022). gp130/STAT3 signaling is required for homeostatic proliferation and anabolism in postnatal growth plate and articular chondrocytes. Communications Biology 5, 1–20.

57. Park, S., Baek, I.J., Ryu, J.H., Chun, C.H., and Jin, E.J. (2022). PPARα-ACOT12 axis is responsible for maintaining cartilage homeostasis through modulating de novo lipogenesis. Nat. Commun. 13. 10.1038/s41467-021-27738-y.

58. Solomon, L.A., Bérubé, N.G., and Beier, F. (2008). Transcriptional regulators of chondrocyte hypertrophy. Birth Defects Res. C Embryo Today 84. 10.1002/bdrc.20124.

59. Laurie, L.E., Kokubo, H., Nakamura, M., Saga, Y., and Funato, N. (2016). The Transcription Factor Hand1 Is Involved In Runx2-Ihh-Regulated Endochondral Ossification. PLoS One 11. 10.1371/journal.pone.0150263.

60. Angerer, H., Krusch, D., Deutsch, A., Gruber, G., Setznagl, D., Kremser, M.L., Leithner, A., Stradner, M.H., and Graninger, W.B. (2012). Potential role of nuclear orphan receptors NR4A1 and NR4A3 in human chondrocytes. Osteoarthritis Cartilage 20, S140.

61. Ghoul-Mazgar, S., Hotton, D., Lézot, F., Blin-Wakkach, C., Asselin, A., Sautier, J.-M., and Berdal, A. (2005). Expression pattern of Dlx3 during cell differentiation in mineralized tissues. Bone 37, 799–809.

62. Song, Z., Lian, X., Wang, Y., Xiang, Y., and Li, G. (2017). KLF15 regulates in vitro chondrogenic differentiation of human mesenchymal stem cells by targeting SOX9. Biochem. Biophys. Res. Commun. 493. 10.1016/j.bbrc.2017.09.078.

63. van der Windt, A.E., Haak, E., Das, R.H.J., Kops, N., Welting, T.J.M., Caron, M.M.J., van Til, N.P., Verhaar, J.A.N., Weinans, H., and Jahr, H. (2010). Physiological tonicity improves human chondrogenic marker expression through nuclear factor of activated T-cells 5 in vitro. Arthritis Res. Ther. 12, R100.

64. Choo, M.-K., Yeo, H., and Zayzafoon, M. (2009). NFATc1 mediates HDAC-dependent transcriptional repression of osteocalcin expression during osteoblast differentiation. Bone 45, 579–589.

65. Cao, Y., Min, J., Zhang, Q., Li, H., and Li, H. (2016). Associations of LBX1 gene and adolescent idiopathic scoliosis susceptibility: a meta-analysis based on 34,626 subjects. BMC Musculoskelet. Disord. 17, 1–10.

66. Kieslinger, M., Folberth, S., Dobreva, G., Dorn, T., Croci, L., Erben, R., Consalez, G.G., and Grosschedl, R. (2005). EBF2 regulates osteoblast-dependent differentiation of osteoclasts. Dev. Cell 9. 10.1016/j.devcel.2005.10.009.

67. Kawai, S., Yamauchi, M., Wakisaka, S., Ooshima, T., and Amano, A. (2007). Zinc-finger transcription factor odd-skipped related 2 is one of the regulators in osteoblast proliferation and bone formation. J. Bone Miner. Res. 22. 10.1359/jbmr.070602.

68. Funato, N., Chapman, S.L., McKee, Funato, H., Morris, J.A., Shelton, J.M., Richardson, J.A., and Yanagisawa, H. (2009). Hand2 controls osteoblast differentiation in the branchial arch by inhibiting DNA binding of Runx2. Development 136. 10.1242/dev.029355.

69. Nakamura, E., Hata, K., Takahata, Y., Kurosaka, H., Abe, M., Abe, T., Kihara, M., Komori, T., Kobayashi, S., Murakami, T., et al. (2021). Zfhx4 regulates endochondral ossification as the transcriptional platform of Osterix in mice. Communications Biology 4, 1–11.

70. Sobolev, V.V., Khashukoeva, A.Z., Evina, O.E., Geppe, N.A., Chebysheva, S.N., Korsunskaya, I.M., Tchepourina, E., and Mezentsev, A. (2022). Role of the Transcription Factor FOSL1 in Organ Development and Tumorigenesis. Int. J. Mol. Sci. 23. 10.3390/ijms23031521.

71. Samee, N., Geoffroy, V., Marty, C., Schiltz, C., Vieux-Rochas, M., Levi, G., and de Vernejoul, M.C. (2008). Dlx5, a positive regulator of osteoblastogenesis, is essential for osteoblast-osteoclast coupling. Am. J. Pathol. 173. 10.2353/ajpath.2008.080243.

72. Veistinen, L.K., Mustonen, T., Hasan, M.R., Takatalo, M., Kobayashi, Y., Kesper, D.A., Vortkamp, A., and Rice, D.P. (2017). Regulation of Calvarial Osteogenesis by Concomitant De-repression of GLI3 and Activation of IHH Targets. Front. Physiol. 8, 1036.

73. Park, R., Madhavaram, S., and Ji, J.D. (2020). The Role of Aryl-Hydrocarbon Receptor (AhR) in Osteoclast Differentiation and Function. Cells 9. 10.3390/cells9102294.

74. Qi, H., Aguiar, D.J., Williams, S.M., La Pean, A., Pan, W., and Verfaillie, C.M. (2003). Identification of genes responsible for osteoblast differentiation from human mesodermal progenitor cells. Proc. Natl. Acad. Sci. U. S. A. 100. 10.1073/pnas.0532693100.

75. Kang, H.C., Chae, J.H., Kim, B.S., Han, S.Y., Kim, S.H., Auh, C.K., Yang, S.I., and Kim, C.G. (2004). Transcription factor CP2 is involved in activating mBMP4 in mouse mesenchymal stem cells. Mol. Cells 17.

76. Franceschi, R.T., and Ge, C. (2017). Control of the Osteoblast Lineage by Mitogen-Activated Protein Kinase Signaling. Current molecular biology reports 3. 10.1007/s40610-017-0059-5.

77. Liu, F., Wang, X., Zheng, B., Li, D., Chen, C., Lee, I.S., Zhong, J., Li, D., and Liu, Y. (2020). USF2 enhances the osteogenic differentiation of PDLCs by promoting ATF4 transcriptional activities. J. Periodontal Res. 55. 10.1111/jre.12689.

78. Bozec, A., Bakiri, L., Jimenez, M., Schinke, T., Amling, M., and Wagner, E.F. (2010). Fra-2/AP-1 controls bone formation by regulating osteoblast differentiation and collagen production. J. Cell Biol. 190, 1093–1106.

79. Huynh, N.P.T., Zhang, B., and Guilak, F. (2019). High-depth transcriptomic profiling reveals the temporal gene signature of human mesenchymal stem cells during chondrogenesis. The FASEB Journal 33, 358.

80. Wu, C.L., Dicks, A., Steward, N., Tang, R., Katz, D.B., Choi, Y.R., and Guilak, F. (2021). Single cell transcriptomic analysis of human pluripotent stem cell chondrogenesis. Nat. Commun. 12. 10.1038/s41467-020-20598-y.

81. Griffiths, R., Woods, S., Cheng, A., Wang, P., Griffiths-Jones, S., Ronshaugen, M., and Kimber, S.J. (2020). The Transcription Factor-microRNA Regulatory Network during hESC-chondrogenesis. Sci. Rep. 10. 10.1038/s41598-020-61734-4.

82. Jääger, K., Islam, S., Zajac, P., Linnarsson, S., and Neuman, T. (2012). RNA-seq analysis reveals different dynamics of differentiation of human dermis- and adipose-derived stromal stem cells. PLoS One 7. 10.1371/journal.pone.0038833.

83. Richard, D., Pregizer, S., Venkatasubramanian, D., Raftery, R.M., Muthuirulan, P., Liu, Z., Capellini, T.D., and Craft, A.M. (2023). Lineage-specific differences and regulatory networks governing human chondrocyte development. Elife 12. 10.7554/eLife.79925.

84. Woods, S., Bates, N., Dunn, S.L., Serracino-Inglott, F., Hardingham, T.E., and Kimber, S.J. (2020). Generation of Human-Induced Pluripotent Stem Cells From Anterior Cruciate Ligament. J. Orthop. Res. 38, 92–104.

85. Young, M.D., Mitchell, T.J., Custers, L., Margaritis, T., Morales-Rodriguez, F., Kwakwa, K., Khabirova, E., Kildisiute, G., Oliver, T.R.W., de Krijger, R.R., et al. (2021). Single cell derived mRNA signals across human kidney tumors. Nat. Commun. 12, 1–19.

86. Robinson, M.D., McCarthy, D.J., and Smyth, G.K. (2010). edgeR: a Bioconductor package for differential expression analysis of digital gene expression data. Bioinformatics 26, 139.

87. Ritchie, M.E., Phipson, B., Wu, D., Hu, Y., Law, C.W., Shi, W., and Smyth, G.K. (2015). limma powers differential expression analyses for RNA-sequencing and microarray studies. Nucleic Acids Res. 43. 10.1093/nar/gkv007.

88. Gong, M., Liang, T., Zhang, H., Chen, S., Hu, Y., Zhou, J., Zhang, X., Zhang, W., Geng, X., and Zou, X. (2018). Gene expression profiling: identification of gene expression in human MSC chondrogenic differentiation. Am. J. Transl. Res. 10, 3555.

89. Dao, D.Y., Jonason, J.H., Zhang, Y., Hsu, W., Chen, D., Hilton, M.J., and O’Keefe, R.J. (2012). Cartilage-specific β-catenin signaling regulates chondrocyte maturation, generation of ossification centers, and perichondrial bone formation during skeletal development. J. Bone Miner. Res. 27. 10.1002/jbmr.1639.

90. Sumanaweera, D., Suo, C., Cujba, A.-M., Muraro, D., Dann, E., Polanski, K., Steemers, A.S., Lee, W., Oliver, A.J., Park, J.-E., et al. (2023). Gene-level alignment of single cell trajectories informs the progression of in vitro T cell differentiation. bioRxiv, 2023.03.08.531713. 10.1101/2023.03.08.531713.

91. Teixeira, C.C., Liu, Y., Thant, L.M., Pang, J., Palmer, G., and Alikhani, M. (2010). Foxo1, a novel regulator of osteoblast differentiation and skeletogenesis. J. Biol. Chem. 285. 10.1074/jbc.M109.079962.

92. Woods, S., Humphreys, P.A., Bates, N., Richardson, S.A., Kuba, S.Y., Brooks, I.R., Cain, S.A., and Kimber, S.J. (2021). Regulation of TGFβ Signalling by TRPV4 in Chondrocytes. Cells 10. 10.3390/cells10040726.

93. Zawel, L., Dai, J.L., Buckhaults, P., Zhou, S., Kinzler, K.W., Vogelstein, B., and Kern, S.E. (1998). Human Smad3 and Smad4 are sequence-specific transcription activators. Mol. Cell 1. 10.1016/s1097-2765(00)80061-1.

94. Humphreys, P.A., Woods, S., Smith, C.A., Bates, N., Cain, S.A., Lucas, R., and Kimber, S.J. (2020). Optogenetic Control of the BMP Signaling Pathway. ACS Synth. Biol. 9, 3067–3078.

95. Korchynskyi, O., and ten Dijke, P. (2002). Identification and functional characterization of distinct critically important bone morphogenetic protein-specific response elements in the Id1 promoter. J. Biol. Chem. 277, 4883–4891.

96. Du, M., Wu, B., Fan, S., Liu, Y., Ma, X., and Fu, X. (2020). SNHG14 induces osteogenic differentiation of human stromal (mesenchymal) stem cells in vitro by downregulating miR-2861. BMC Musculoskelet. Disord. 21. 10.1186/s12891-020-03506-9.

97. LncRNA KCNQ1OT1 promotes osteogenic differentiation via miR-205-5p/RICTOR axis (2022). Exp. Cell Res. 415, 113119.

98. Jensen, E.D., Niu, L., Caretti, G., Nicol, S.M., Teplyuk, N., Stein, G.S., Sartorelli, V., van Wijnen, A.J., Fuller-Pace, F.V., and Westendorf, J.J. (2008). p68 (Ddx5) interacts with Runx2 and regulates osteoblast differentiation. J. Cell. Biochem. 103, 1438–1451.

99. Hjorten, R., Hansen, U., Underwood, R.A., Telfer, H.E., Fernandes, R.J., Krakow, D., Sebald, E., Wachsmann-Hogiu, S., Bruckner, P., Jacquet, R., et al. (2007). Type XXVII collagen at the transition of cartilage to bone during skeletogenesis. Bone 41, 535.

100. Yang, L., Tsang, K.Y., Tang, H.C., Chan, D., and Cheah, K.S.E. (2014). Hypertrophic chondrocytes can become osteoblasts and osteocytes in endochondral bone formation. Proc. Natl. Acad. Sci. U. S. A. 111, 12097–12102.

101. Hirsch, C., and Schildknecht, S. (2019). In Vitro Research Reproducibility: Keeping Up High Standards. Front. Pharmacol. 10. 10.3389/fphar.2019.01484.

102. Riss, T.L., Moravec, R.A., Duellman, S.J., and Niles, A.L. (2021). Treating Cells as Reagents to Design Reproducible Assays. SLAS discovery : advancing life sciences R & D 26. 10.1177/24725552211039754.

103. Yamini Krishnan, A.J.G. (2018). Cartilage Diseases. Matrix Biol. 71–72, 51.

104. Inacio, M.C.S., Paxton, E.W., Graves, S.E., Namba, R.S., and Nemes, S. (2017). Projected increase in total knee arthroplasty in the United States – an alternative projection model. Osteoarthritis and Cartilage 25, 1797–1803. 10.1016/j.joca.2017.07.022.

105. Singh, J.A. (2011). Epidemiology of Knee and Hip Arthroplasty: A Systematic Review§. The Open Orthopaedics Journal 5, 80–85. 10.2174/1874325001105010080.

106. McCaskie, A.W. (2015). From needle to knife. Bone Joint J. 97-B, 1–2.

## Methods References

1. Ye, J., Bates, N., Soteriou, D., Grady, L., Edmond, C., Ross, A., Kerby, A., Lewis, P.A., Adeniyi, T., Wright, R., et al. (2017). High quality clinical grade human embryonic stem cell lines derived from fresh discarded embryos. Stem Cell Res. Ther. 8. 10.1186/s13287-017-0561-y.

2. Dobin, A., Davis, C.A., Schlesinger, F., Drenkow, J., Zaleski, C., Jha, S., Batut, P., Chaisson, M., and Gingeras, T.R. (2013). STAR: ultrafast universal RNA-seq aligner. Bioinformatics 29, 15–21.

3. Frankish, A., Diekhans, M., Ferreira, A.-M., Johnson, R., Jungreis, I., Loveland, J., Mudge, J.M., Sisu, C., Wright, J., Armstrong, J., et al. (2019). GENCODE reference annotation for the human and mouse genomes. Nucleic Acids Res. 47, D766–D773.

4. Robinson, J.T., Thorvaldsdóttir, H., Winckler, W., Guttman, M., Lander, E.S., Getz, G., and Mesirov, J.P. (2011). Integrative genomics viewer. Nature Biotechnology 29, 24–26. 10.1038/nbt.1754.

5. Li, B., and Dewey, C.N. (2011). RSEM: accurate transcript quantification from RNA-Seq data with or without a reference genome. BMC Bioinformatics 12. 10.1186/1471-2105-12-323.

6. Robinson, M.D., McCarthy, D.J., and Smyth, G.K. (2010). edgeR: a Bioconductor package for differential expression analysis of digital gene expression data. Bioinformatics 26, 139–140.

7. Gu, Z., Eils, R., and Schlesner, M. (2016). Complex heatmaps reveal patterns and correlations in multidimensional genomic data. Bioinformatics 32, 2847–2849.

8. Kaminow, B., Yunusov, D., and Dobin, A. (2021). STARsolo: accurate, fast and versatile mapping/quantification of single-cell and single-nucleus RNA-seq data. bioRxiv, 2021.05.05.442755. 10.1101/2021.05.05.442755.

9. Lun, A.T.L., Riesenfeld, S., Andrews, T., Dao, T.P., Gomes, T., participants in the 1st Human Cell Atlas Jamboree, and Marioni, J.C. (2019). EmptyDrops: distinguishing cells from empty droplets in droplet-based single-cell RNA sequencing data. Genome Biol. 20, 63.

10. Young, M.D., and Behjati, S. (2020). SoupX removes ambient RNA contamination from droplet-based single-cell RNA sequencing data. Gigascience 9. 10.1093/gigascience/giaa151.

11. Wolf, F.A., Angerer, P., and Theis, F.J. (2018). SCANPY: large-scale single-cell gene expression data analysis. Genome Biol. 19, 15.

12. Polański, K., Young, M.D., Miao, Z., Meyer, K.B., Teichmann, S.A., and Park, J.-E. (2020). BBKNN: fast batch alignment of single cell transcriptomes. Bioinformatics 36, 964–965.

13. GitHub - cellgeni/sceasy: A package to help convert different single-cell data formats to each other GitHub. https://github.com/cellgeni/sceasy.

14. Wickham, H. (2009). ggplot2: Elegant Graphics for Data Analysis (Springer Science & Business Media).

15. Wolock, S.L., Lopez, R., and Klein, A.M. (2019). Scrublet: Computational Identification of Cell Doublets in Single-Cell Transcriptomic Data. Cell Syst 8, 281–291.e9.

16. Eli Bingham Uber AI Labs, Uber Technologies, Inc., San Francisco, CA, Jonathan P. Chen Uber AI Labs, Uber Technologies, Inc., San Francisco, CA, Martin Jankowiak Uber AI Labs, Uber Technologies, Inc., San Francisco, CA, Fritz Obermeyer Uber AI Labs, Uber Technologies, Inc., San Francisco, CA, Neeraj Pradhan Uber AI Labs, Uber Technologies, Inc., San Francisco, CA, Theofanis Karaletsos Uber AI Labs, Uber Technologies, Inc., San Francisco, CA, Rohit Singh Uber AI Labs, Uber Technologies, Inc., San Francisco, CA, Paul Szerlip Uber AI Labs, Uber Technologies, Inc., San Francisco, CA, Paul Horsfall Uber AI Labs, San Francisco, CA and Stanford University, Stanford, CA, and Noah D. Goodman Uber AI Labs, San Francisco, CA and Stanford University, Stanford, CA (2019). Pyro. J. Mach. Learn. Res. 10.5555/3322706.3322734.

17. Dann, E., Henderson, N.C., Teichmann, S.A., Morgan, M.D., and Marioni, J.C. (2021). Differential abundance testing on single-cell data using k-nearest neighbor graphs. Nat. Biotechnol. 40, 245–253.

18. Garcia-Alonso, L., Holland, C.H., Ibrahim, M.M., Turei, D., and Saez-Rodriguez, J. (2019). Benchmark and integration of resources for the estimation of human transcription factor activities. Genome Res. 29, 1363–1375.

19. Holland, C.H., Tanevski, J., Perales-Patón, J., Gleixner, J., Kumar, M.P., Mereu, E., Joughin, B.A., Stegle, O., Lauffenburger, D.A., Heyn, H., et al. (2020). Robustness and applicability of transcription factor and pathway analysis tools on single-cell RNA-seq data. Genome Biol. 21, 36.

20. Alvarez, M.J., Shen, Y., Giorgi, F.M., Lachmann, A., Ding, B.B., Ye, B.H., and Califano, A. (2016). Functional characterization of somatic mutations in cancer using network-based inference of protein activity. Nat. Genet. 48, 838–847.

21. Young, M.D., Mitchell, T.J., Custers, L., Margaritis, T., Morales, F., Kwakwa, K., Khabirova, E., Kildisiute, G., Oliver, T.R.W., de Krijger, R.R., et al. (2020). Single cell derived mRNA signals across human kidney tumors. bioRxiv, 2020.03.19.998815. 10.1101/2020.03.19.998815.

22. Sumanaweera, D., Suo, C., Cujba, A.-M., Muraro, D., Dann, E., Polanski, K., Steemers, A.S., Lee, W., Oliver, A.J., Park, J.-E., et al. (2023). Gene-level alignment of single cell trajectories informs the progression of in vitro T cell differentiation. bioRxiv, 2023.03.08.531713. 10.1101/2023.03.08.531713.

23. Wallace, C.S., and Freeman, P.R. (1987). Estimation and Inference by Compact Coding. J. R. Stat. Soc. Series B Stat. Methodol. 49, 240–252.

24. Navas-Palencia, G. (2020). Optimal binning: mathematical programming formulation.

25. Lambert, S.A., Jolma, A., Campitelli, L.F., Das, P.K., Yin, Y., Albu, M., Chen, X., Taipale, J., Hughes, T.R., and Weirauch, M.T. (2018). The Human Transcription Factors. Cell 175, 598–599.

26. Kanehisa, M., Furumichi, M., Sato, Y., Kawashima, M., and Ishiguro-Watanabe, M. (2023). KEGG for taxonomy-based analysis of pathways and genomes. Nucleic Acids Res. 51. 10.1093/nar/gkac963.

27. Lee, H., Marco Salas, S., Gyllborg, D., and Nilsson, M. (2022). Direct RNA targeted in situ sequencing for transcriptomic profiling in tissue. Sci. Rep. 12, 1–9.

28. Gataric, M., Park, J.S., Li, T., Vaskivskyi, V., Svedlund, J., Strell, C., Roberts, K., Nilsson, M., Yates, L.R., Bayraktar, O., et al. (2021). PoSTcode: Probabilistic image-based spatial transcriptomics decoder. bioRxiv, 2021.10.12.464086. 10.1101/2021.10.12.464086.

29. Vaskivskyi Vasyl (2022). MicroAligner.

30. Allan, D.B., Caswell, T., Keim, N.C., van der Wel, C.M., and Verweij, R.W. (2023). soft-matter/trackpy: v0.6.1 (Zenodo) 10.5281/ZENODO.7670439.

31. Stringer, C., Wang, T., Michaelos, M., and Pachitariu, M. (2020). Cellpose: a generalist algorithm for cellular segmentation. Nat. Methods 18, 100–106.

32. Sean Gillies (2007). Shapely: manipulation and analysis of geometric objects. https://shapely.readthedocs.io/en/stable/index.html.

